# Genotypic and environmental effects on seed coat patterning and nutritional composition in common bean (*Phaseolus vulgaris* L.)

**DOI:** 10.64898/2026.04.13.718301

**Authors:** Tayah M. Bolt, Austin Cole, Rajdeep Bains, Li Tian, Travis A. Parker, Paul Gepts, Antonia Palkovic, Gail M. Bornhorst, Christine H. Diepenbrock

## Abstract

Common bean (*Phaseolus vulgaris* L.) is the leading grain legume consumed directly by humans and a primary source of nutrients in many communities. This study utilized common bean genotypes with diverse seed coat phenotypes to investigate genotypic and environmental effects on pigmented seed coat area and seed macronutrient (protein, starch, fat, ash, moisture), anti-nutrient (phytate), and mineral nutrient (iron, zinc, calcium, phosphorus, magnesium, potassium, sodium) profiles. Recombinant inbred lines (RILs) that comprise six phenotypic classes for seed coat patterning and nine commercial cultivars were field-evaluated for multiple years across inland, coastal, and intermountain environments in California. A custom near-infrared spectroscopy calibration improved macronutrient prediction accuracy relative to a pre-existing calibration. Environmental effects on macronutrients were pronounced; the 2022 coastal growing environment was the most distinct, characterized by significantly higher starch and moisture content and significantly lower protein content in the RILs relative to any other environments. Across growing years in the RILs, greater consistency was observed at the inland site, where only protein was significantly different; all macronutrient traits significantly differed within the intermountain site. Certain commercial cultivars largely maintained their relative rank for protein content across environments, indicating consistency of genotypic performance, and Black Nightfall ranked among the highest for iron, zinc, phosphorus, and magnesium. Percent pigmented seed coat area was significantly negatively correlated with both calcium and magnesium concentrations. These results underscore the importance of genotype-by-environment field trials for seed coat patterning, seed nutritional composition, and their interplay, to support breeding of common bean among other grain legumes.

**Highlights:**

- Custom near-infrared spectroscopy (NIRS) calibration improved prediction accuracies
- Environmental effects significantly influenced common bean macronutrient composition
- Certain cultivars ranked consistently for macronutrient traits across environments
- Seed coat pattern was significantly associated with mineral nutrient concentrations

## 1 Introduction

Common bean (*Phaseolus vulgaris* L.) is the leading grain legume consumed directly by humans, and is grown globally with major production in the Americas and Asia alongside cultivation in Africa and Mediterranean Europe ^1–3^. For many communities in sub-Saharan Africa and Latin America, including countries such as Rwanda, Kenya, Uganda, Mexico, and Brazil, common bean seed (or grain) serves as a primary source of protein ^1,4^. Common bean also typically contains high grain mineral content, providing a dietary source of iron, zinc, phosphorus, copper, and aluminum ^1,5^. The nutritional value and diverse culinary history of common bean makes it extremely culturally important as a crop and dietary constituent ^1,6–9^. Common bean includes many commercially relevant market classes that differ in their seed coat color (e.g., kidney, black, navy, Great Northern, pink, and yellow beans) and seed coat patterning (e.g., pinto, Flor de Mayo, red mottled, and cranberry beans) ^10^. As consumers and growers often exhibit strong preferences for dry beans with specific seed coat colorations/patterns, a diversity of seed types can be observed in common bean, and consumers are willing to pay a premium for specialty dry beans (i.e., cultivars grown for specific quality traits such as unique colorations/patterns, sensory attributes, nutritional profiles, or regionally adapted agronomic characteristics) ^11–14^.

Environmental effects are important when considering plant breeding for both improved macronutrient and micronutrient profiles and preferred seed coat patterns ^15^. For example, in common bean it has been shown that the environment has a major effect on seed zinc and iron concentrations as well as pigmented seed coat area ^16,17^. These traits are often highly sensitive to environmental variables such as elevation, weather conditions (e.g. rainfall and temperature), and soil properties/amendments (e.g., pH and nitrogen fertilization), which can influence nutrient uptake, metabolic pathways, and overall plant physiology ^18,19^. Thus, it is essential to evaluate genotypes in multi-environment trials to accurately identify high-performing genotypes for use in breeding programs working on nutritional quality alongside priority agronomic traits for production by farmers in the target population of environments (TPE). In California, this TPE for common bean spans coastal, inland, and intermountain geographies ^20–22^. Favorable and stable quality is also important for the continued contributions of dry beans as a low-input rotation crop in farming systems in these geographies ^1,23^. Despite the agronomic and nutritional importance of common bean, genotype by environment (GxE) studies remain geographically concentrated. A recent systematic review by Karavidas et al.^24^ of 228 published studies focused on agronomic practices for yield and quality in common bean reported that the majority of research production sites have been in Asia and South America, underscoring the need for broader multi-environment evaluations. While some GxE studies for yield have been conducted in the United States ^25,26^, the examination of quality traits across diverse growing regions is critical for identifying genotypes with favorable and stable performance for seed coat patterning, macronutrient, mineral nutrient, and/or anti-nutrient traits.

Malnutrition, including both undernutrition and micronutrient deficiencies, globally affected over 194 million children under the age of five years as of 2020 ^27^, and micronutrient deficiencies, or “hidden hunger,” are prevalent in regions where common bean is a dietary staple ^28,29^. Additionally, protein–energy malnutrition, the most widespread deficiency disorder in many developing countries, primarily affects young children and leads to high mortality in severe cases and lasting developmental impairments in milder forms ^30^. Pressing concerns of growing population size, current and projected environmental stresses on crop performance, market dynamics, and the persistence of malnutrition underscore the need for nutritionally dense foods and the need to improve our understanding of how crop quality profiles vary across crop genotypes and production environments. While plant breeding efforts have largely been focused on increasing crop yield and usability rather than nutritional quality traits ^31^, more focus has been brought to nutritional quality with the rise of biofortification, which is the improvement of crop nutritional quality through plant breeding and/or agronomic approaches ^32,33^. Micronutrients and anti-nutrients have critical roles in addressing micronutrient deficiencies, and iron has been a key target for biofortification efforts in common bean ^34,35^. Anti-nutrients can significantly affect the bioavailability (availability for utilization in the human body ^35^) of both macro- and micronutrients, making their mitigation important for improving the effectiveness of biofortification strategies ^29,36^. As an example of a major anti-nutrient in common bean among other staples, phytate is the primary storage form of seed phosphorus but can also bind proteins as well as iron, magnesium, and zinc, limiting their bioavailability ^37,38^. Bouis and Welch^34^ report that overall, seeds and grains of staple crops provide only low levels of bioavailable iron and zinc, with roughly 5% of the total seed iron and 25% of the total seed zinc in common bean estimated to be available for absorption due to the presence of anti-nutrients such as phytate. Low-phytate beans have been previously developed; however, they have also been associated with adverse gastrointestinal symptoms, likely due to the presence of phytohemagglutinin-L, a lectin that remained in the beans after cooking ^38–40^. Thus, it is of interest to decrease phytate levels while being mindful of potential tradeoffs in common bean breeding efforts ^41^. Additional anti-nutrients present in common bean include specific phenolic compounds such as myricetin, myricetin 3-glucoside, quercetin, and quercetin 3-glucoside, which have been found to inhibit iron bioavailability, while other phenolics have been found to promote iron uptake ^41–43^. As a result, lowering the concentration of anti-nutrients could be used as a strategy to enhance the nutritional value of common bean by increasing the bioavailability of existing macro- and micronutrients ^34,44,45^.

In this study, to investigate genotypic and environmental effects on seed coat patterning and nutritional composition, we evaluated a total of 38 common bean genotypes in small-plot trials from 2022 to 2024, across three subregions within California with varying day and night temperatures, for a total of seven growing environments. These genotypes included recombinant inbred lines (RILs) representing six phenotypic classes for seed coat patterning, their parents (Black Nightfall and Orca), and seven commercially relevant cultivars (Figure 1). Samples from these plots were analyzed for pigmented seed coat area and macronutrient profiles (protein, starch, ash, and moisture content as a percentage of seed composition). In the three trials (in each of 2022-2024) conducted in Davis, CA, phytate and mineral nutrient levels (e.g., for iron, zinc, calcium, phosphorus, and magnesium) were also measured. We hypothesized that (i) pigmented seed coat area and nutritional traits are significantly affected by genotype, environment, and their interaction, and (ii) pigmented seed coat area is associated with variation in seed nutritional composition. A custom near-infrared spectroscopy calibration was built for use on this sample set; linear mixed-effects modeling was used to dissect the main and interaction effects of genotype and environment on the aforementioned traits; and relationships among these traits were assessed in RILs and commercial cultivars. This collective information can help to improve our foundational understanding of the genetic and environmental basis of nutritional quality in common bean and build towards the development of predictive models for application in breeding pipelines.

**Figure 1.**
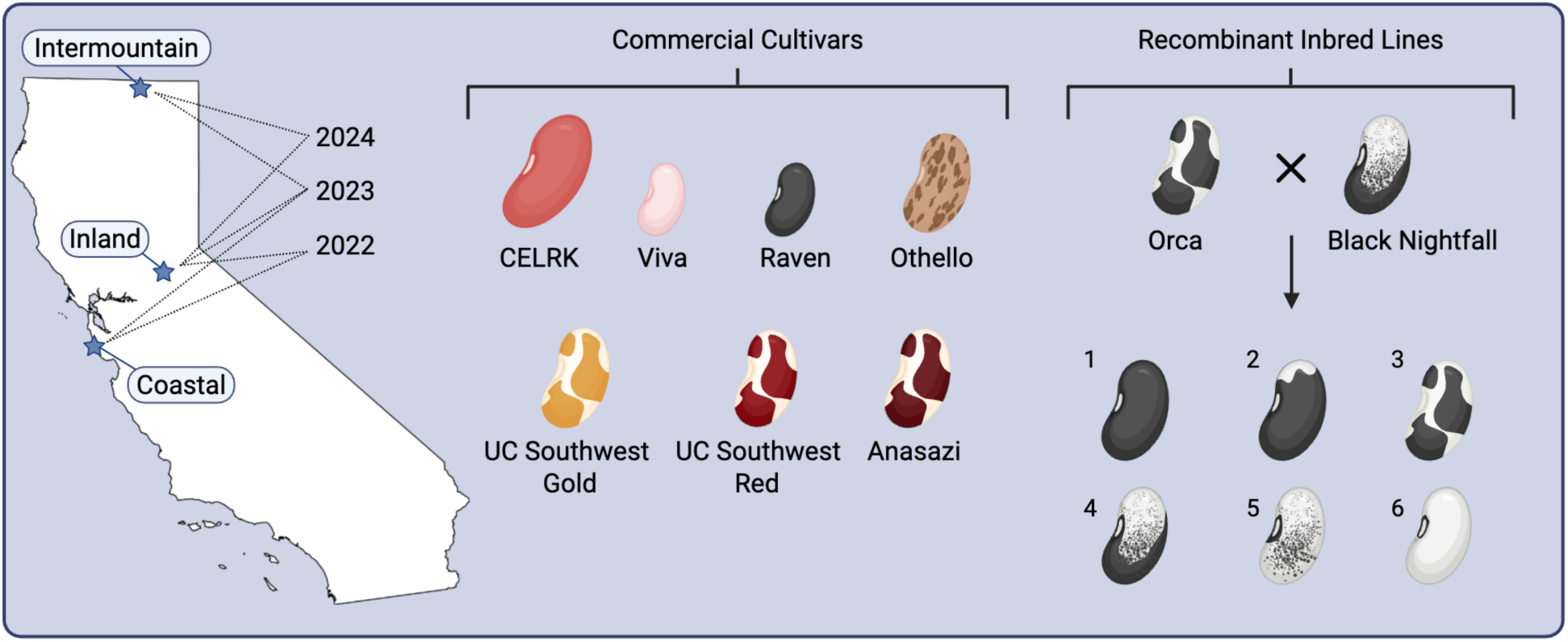
Seed coat phenotypes and growing locations across California (2022–2024) for commercial cultivars and recombinant inbred lines. The intermountain, inland, and coastal locations are represented by the top, middle, and bottom stars, respectively. Seed coat coloration and patterning, as well as approximate seed size, are depicted for each commercial cultivar and RIL phenotypic class. Acronyms: CELRK, California Early Light Red Kidney; UC, University of California.

## 2 Materials and Methods

### 2.1 Material Selection and Growing Conditions

This study utilized common bean genotypes with varying seed coat colorations and patterning. These genotypes included recombinant inbred lines (RILs) of six phenotypic classes^46^, their parents (Black Nightfall [W6 51267] and Orca [PI 632344]^47^), and seven commercially relevant cultivars, namely Anasazi (PI 577705; Adobe Milling)^48^, University of California Southwest Red (UCSW Red) (PI 693472)^48^, University of California Southwest Gold (UCSW Gold) (PI 693470)^49^, and one variety from each of the following major market classes of dry bean: pinto (Othello [P1 578268])^50^, black (Raven [P1 57807])^51^, pink (Viva [PI 549940])^52^, and kidney (California Early Light Red Kidney; CELRK)^53^ (Figure 1; Table S1). The phenotypic classes for RILs are based on genotypic classes from a three-gene model between the *P*, *T*, and *Bip* genes.

These genes have been mapped to *PvTT8*, *PvTTG1*, and *PvMYC1*, respectively, which are predicted to be members of MYB-bHLH-WD40 complexes ^16,54^. Eight genotypic classes give rise to six unique phenotypic classes (Figure 1), as *Bip* is hypostatic to *T*. A total of 29 RILs were utilized in this study, with selection stratified by phenotypic class (based on the number of RILs per phenotypic class to be evaluated in each environment, as described below; Table S1) and based on seed availability. Black Nightfall and Orca are members of phenotypic classes 4 and 3, respectively. Anasazi and UC Southwest Red and Gold also exhibit seed coat patterns similar to those of phenotypic class 3 (Figure 1).

All seed samples analyzed in this study were grown in small-plot trials across California in 2022, 2023, and 2024. The number of genotypes (18 to 33) and biological replicate plots (two or three per genotype) planted each year was dependent on the number of plots available for use in each field as well as RIL seed stock availability (Table S1). In 2022, seed was grown in both a coastal location (San Gregorio, California) and an inland location (Davis, California). In 2023, seed was grown in a coastal location (Santa Cruz, California), an inland location (Davis, California), and an intermountain location (Tulelake, California). In 2024, seed was grown in both an inland location (Davis, California) and an intermountain location (Tulelake, California). Four RILs per phenotypic class were evaluated in 2022, and three RILs per phenotypic class were evaluated in 2023 and 2024, with the exception that only two RILs per phenotypic class were evaluated in 2023 in Santa Cruz due to space limitations (Table S1).

Seed produced in San Gregorio, California, in 2022 was from an organic on-farm trial planted on June 14 using two-row plots that were 20 feet long by 5 feet wide. Two plots for each of 33 genotypes (Table S1) were grown in a randomized complete block design, with flanking borders of continuously planted UC Southwest Gold. Overhead irrigation was used after planting to promote successful emergence, and the beans were then dry-farmed for the remainder of the season. Plants were pulled from the ground on September 23 to complete senescence/drydown in the field, and seed pods were harvested from the field on September 29. Samples were threshed via stationary thresher one day after harvest. All samples were stored at -80°C on the day that they were threshed.

Seed produced in Santa Cruz, California, in 2023 was from an organic farm trial at the UC Santa Cruz Center for Agroecology Farm using two-row plots that were 12 feet long by 6 feet wide, and 91.02 kg of compost per ha was applied to the field. Seed for 18 genotypes (Table S1) was planted on June 6, pulled from the ground on October 9, and seed pods were harvested from the field on October 11. Two plots for each genotype were planted utilizing a randomized complete block design, and an additional third plot for four of the 18 genotypes (Orca, Black Nightfall, UCSW Gold, and UCSW Red) was planted due to limited additional available space. The field was irrigated during planting and then dry farmed. Samples were threshed via stationary thresher one day after harvest. All samples were stored at -80°C on the day that they were threshed.

For seed produced in Tulelake, California, in 2023, seed was planted on May 26, cut and windrowed on Sept. 14/15, and harvested/threshed in the field on September 20. For seed produced in Tulelake, California, in 2024, seed was planted on May 31, cut and windrowed on September 30, and harvested/threshed in the field on October 3. All years were sprinkler-irrigated at this planting location. In 2023, two-row plots that were 20 feet long by 5 feet wide were used, and 112.10 kg of nitrogen per ha was applied to the field. In 2024, three-row plots, 10 feet long by 5 feet wide, were used, and 123.31 kg of nitrogen per ha was applied. Two plots per genotype were grown in 2023, and three plots per genotype were grown in 2024, with both years utilizing a randomized complete block design for 27 genotypes and flanking borders of continuously planted UC Southwest Gold (Table S1). All samples were stored at -80°C on the day that they were threshed.

For seed produced in Davis, California, in 2022, 33 genotypes (Table S1) were planted on May 25, cut and windrowed on August 17, and harvested/threshed in the field on September 6-7. For seed produced in Davis, California, in 2023, 24 genotypes (Table S1) were planted on May 24, cut and windrowed on September 5, and harvested/threshed in the field on September 12. For seed produced in Davis, California, in 2024, 27 genotypes (Table S1) were planted on May 23, cut and windrowed on September 16, and harvested/threshed in the field on September 30. All years were irrigated via buried drip at this planting location, and all years used two-row plots that were 10 feet long by 5 feet wide. In 2022, 2023, and 2024, the following amounts of nitrogen were applied to the field: 30.27, 84.08 (due to weed pressure), and 31.39 kg per ha, respectively. Three plots for each genotype were grown in a randomized complete block design in both 2022 and 2023. In 2024, three plots were planted per RIL, while two plots were planted per commercial cultivar. All samples were stored at -80°C on the day that they were threshed. Additionally, for this location, soil samples were taken in triplicate each year from the field post-harvest on October 21, 2022; December 1, 2023; and October 24, 2024, respectively.

### 2.2 Whole Seed Imaging

For the quantification of percent pigmented seed coat area, whole beans were thawed and imaged against a blue contrast background and analyzed in ImageJ as described by Parker et al. ^16^.

Images were taken using a Brother (Nagoya, Japan) DCP-7065DN scanner, then processed using custom macros that were developed in ImageJ (source code available at https://github.com/TravisParker91/Seed-color). This image analysis produced raw data for both the percentage of the image that was pigmented seed coat and the percentage of the image that was seed. To calculate the proportion of the seed coat that was pigmented, the first trait was divided by the second trait, then multiplied by 100 to represent the percent pigmented seed coat area.

### 2.3 At-Harvest Sample Preparation

Post-imaging, seed grown in 2022 was pooled, where samples from the two or three plots planted for each genotype at each location were combined due to freezer space limitations. Prior to sample pooling, a subset of seven genotypes from Davis and six genotypes from San Gregorio with sufficient seed for by-plot analysis were analyzed for macronutrient traits (protein, starch, fat, crude fiber, ash, and moisture) (Supplemental Table S2). As the standard deviation between plots was low for all traits, all further analyses were conducted on the pooled samples for 2022, while 2023 and 2024 were analyzed on a by-plot basis. Grain yield (g) was obtained for the pooled San Gregorio 2022 samples and the Davis 2024 by-plot samples, but quantification of grain yield was not feasible for the remaining growing environments.

Once imaged, whole raw seeds were ground to a fine powder using an IKA tube mill (Tube Mill control S001, Wilmington, NC, USA) with multi-use milling tubes (i.e., grinding chambers; IKA MMT 40.1, Wilmington, NC, USA). Approximately 40 g of sample was split into two subsets (to avoid overloading the grinding chamber), then ground at 25,000 rpm for 78 seconds in 10-second intervals (the final interval was only 8 seconds in duration), and the two ground subsets were then recombined. Ground samples were stored at -80°C until further analysis.

### 2.4 Macronutrient Analysis

#### 2.4.1 Near-infrared Spectroscopy

Ground samples were scanned via near-infrared spectroscopy (NIRS), using a pre-existing Vegetal Protein Meals (VPM) calibration available from the NIRS instrument manufacturer. The instrument used was a FOSS DS2500, which provides reflectance data from 400 to 2500 nm at each 0.5 nm step. The VPM calibration predicts percent of protein, starch, fat, ash (total mineral), crude fiber, and moisture from spectral data and was built on samples representing multiple plant protein meals (namely from soybean and other oilseeds). To validate the predicted values generated from the NIR spectra via this calibration and to inform the training of a custom calibration, a subset of the samples (selected via Kennard-Stone algorithm) also underwent chemical analysis as described in section 2.4.2. The data from those analyses were then used to train a custom calibration (described in section 2.7) to predict macronutrient and mineral nutrient trait values from the NIR spectra. For macronutrients, results were then compared with predictions obtained from the pre-existing VPM calibration.

#### 2.4.2 Chemical Analyses

The chemical analyses described in this section were conducted on a selected subset of the NIRS-scanned samples. For the 2022 samples, all pooled samples were selected for a total of 66 samples. Additionally, four UCSW Gold samples from the 2022 San Gregorio border were selected: two from the east border and two from the west. For the 2023 and 2024 samples, NIRS spectra were analyzed via the Kennard-Stone algorithm in RStudio^55^ (using the *kenStone()* function of the R package prospectr, with the metric argument set to ‘mahal’ for Mahalanobis distance) to identify the subset of 30 by-plot samples from each 2023 and 2024 location to use for chemical analyses, resulting in 150 samples ^56^. In total, of the 400 scanned samples, 220 were selected for chemical validation.

Total starch for the ground dry bean samples was determined using a Megazyme Total Starch Assay Kit (AA/AMG, PC: K-TSTA-100A, Wicklow, Ireland) following the AOAC Official Method 996.11,^57^ with minimal adaptation. Specifically, rather than measuring individual reactions in a glass conical tube, the final reaction values were measured at 510 nm by loading 225 µL in triplicate into a 96-well plate to be read on a multi-mode plate reader.

Ground dry bean samples were also sent to the UC Davis Analytical Lab to determine dry matter, protein, fat, and ash values. Dry matter was measured by oven drying for 3 hours at 105°C. Protein was measured by the AOAC Official Method 990.03,^58^ Protein (Crude) in Animal Feed, via the combustion method. Ash was measured by the AOAC Official Method 942.05,^59^ Ash of Animal Feed. Fat was measured by the AOAC Official Method 2003.05,^60^ Crude Fat in Feeds, Cereal Grains, and Forages. Each trait value was reported as a percentage of seed composition on an “as is” basis (i.e., without having corrected for moisture).

### 2.5 Micronutrient and Anti-nutrient Analysis

As soil samples were only taken in Davis, CA, and beans were grown in Davis, CA, in all three trial years, micronutrients and anti-nutrients were measured only in the aforementioned Davis, CA, subset of samples (all pooled samples in 2022, and the samples selected via the Kennard-Stone algorithm in each of 2023 and 2024).

Phytate content (as a percentage of seed composition) for the ground dry bean samples was determined using a Megazyme Phytic Acid (Phytate)/Total Phosphorus Assay Kit (PC: K-PHYT, Wicklow, Ireland) following the provided standard assay procedure, with minimal adaptation. Specifically, rather than measuring individual reactions in a glass conical tube, the final reaction values were measured at 655 nm by loading 225 µL in triplicate into a 96-well plate to be read on a multi-mode plate reader.

For the assessment of iron, zinc, calcium, phosphorus, magnesium, potassium, and sodium, ground raw bean samples were sent to the UC Davis Interdisciplinary Center for Plasma Mass Spectrometry for analysis via inductively coupled plasma mass spectrometry (ICP-MS).

Prior to ICP-MS analysis, samples underwent CEM MARS6 microwave acid digestion using Xpress vessels. The microwave digestion program was run using a 15 min ramp and 15 min hold at 180°C with ∼100 mg bean samples alongside digestion batch quality control samples and 1.00 mL concentrated trace-metal grade HNO₃. Samples were brought to a final volume of 10.00 mL with Milli-Q water.

Microwave-digested bean samples, along with digestion batch quality control (QC) samples, were analyzed using an Agilent 8900 ICP-MS Triple Quad (Agilent Technologies, Santa Clara, CA, USA). The instrument was equipped with nickel interface cones and an x-Lens and was operated in MS/MS helium mode using a three-point peak pattern, with three replicates per injection, and 50 sweeps per replicate. Samples and calibration standards were introduced via an Agilent SPS 4 Autosampler, integrated peristaltic pump, and a MicroMist nebulizer, to generate an aerosol that passed through a Scott-type spray chamber maintained at 2°C and into a plasma operated at 1550 W. Helium was used as the collision cell gas to mitigate polyatomic interferences.

External calibration standards were prepared from single-element phosphorus and zinc standards (Inorganic Ventures) and Calibration Standard 3 (SPEX CertiPrep). A custom internal standard solution containing scandium, germanium, yttrium, indium, and bismuth was prepared from single-element standards (Inorganic Ventures). All standards and samples were prepared using concentrated trace-metal grade HNO₃ and Milli-Q water, with final solutions containing 10% (v/v) trace-metal grade HNO₃.

Digestion batch QC samples included method blanks (Milli-Q water), Laboratory Control Standard A (custom multi-element standard; Inorganic Ventures), and Laboratory Control Standard B (NIST 1573a Tomato Leaves). These QC samples were analyzed alongside digested bean samples. For initial calibration and blank verification, matrix-matched NIST 1643f, an independent-source single-element phosphorus standard (Inorganic Ventures), and 10% (v/v) trace-metal grade HNO₃ were analyzed. Continuing calibration verification standards were prepared from single-element phosphorus and zinc standards (Inorganic Ventures) and Calibration Standard 3 (SPEX CertiPrep), with 10% (v/v) trace-metal grade HNO₃ serving as the continuing calibration blanks.

Raw data were processed using MassHunter ICP-MS software version 4.6 (Agilent). Data reduction included application of a correction equation to account for Sr²⁺ double-charged interference on calcium.

### 2.6 Environmental Analysis

Soil samples were sent to the UC Davis Analytical Lab to determine pH. Additionally, soil samples were sent to the UC Davis Interdisciplinary Center for Plasma Mass Spectrometry for analysis via ICP-MS for the quantification of iron, zinc, calcium, phosphorus, magnesium, potassium, and sodium. Prior to ICP-MS, samples underwent CEM MARS6 microwave acid digestion as described in section 2.5, with the minor adjustments of ∼50 mg of soil sample being used, and Laboratory Control Standard B was NIST 2711a Montana II Soil. ICP-MS analysis was then carried out as described above in section 2.5.

Weather data for each growing environment over the time period of planting to harvest was obtained via the NASA Prediction Of Worldwide Energy Resources Data Access Viewer (Figure S1) ^61^.

### 2.7 NIRS Calibration, Statistical Analysis, and Visualization

The following was conducted using RStudio (RStudio, 2021.09.1, Build 372, © 2009-2021 RStudio, PBC) ^55,62^.

#### 2.7.1 NIRS Custom Calibration

For the custom NIRS calibration, all spectral data was first pre-treated using the standard normal variate (SNV) filter via the *standardNormalVariate()* function of the R package prospectr ^56^. Multiple spectral pre-treatment methods were evaluated, and SNV was selected as it provided the best predictive performance (indicated by the highest correlation between predicted and observed values) and the lowest root mean square error of prediction (RMSEP) in five-fold cross-validation. Extreme outliers were then detected in the data set by the calculation of Mahalanobis distance using the function *mahalanobis()*. Outliers with Mahalanobis distances in the 99.5th percentile threshold were removed, resulting in the removal of two samples from the dataset. The detection of outliers from the chemical analyses data set was performed in the same way, however no samples were removed. Partial least squares regression (via the *plsr()* function) with R package pls was used for five-fold cross-validation within the complete set of data which had both spectral and chemical reference values ^63^. For each trait, the *selectNcomp()* function was used to select the number of principal components to use for the final predictions. The selected number of principal components for each trait was visually confirmed via interactive graphic interface using the RMSEP value from the five-fold cross-validation plotted against principal component number. If the number of components selected was not appropriately selected via the onesigma method (i.e., a lower RMSEP could be obtained using a higher number of principal components), the number of components was overridden and hand selected (Supplemental File S1). The *predict()* function was used to predict new trait values using the SNV pre-treatment and the same optimal number of components identified in five-fold cross-validation. In total, 394 samples received final predictions for protein, total nitrogen, starch, crude fat, ash, and moisture from this custom, PLS-based calibration; these final values were used for further linear mixed-effects modeling and statistical analysis. The predicted (via near-infrared spectroscopy; NIRS) vs. observed (chemical reference) values for protein, starch, crude fat, ash (total mineral), and moisture were then visualized using the *ggplot()* function from the ggplot2 R package ^64^.

#### 2.7.2 Linear Mixed-Effects Modeling, Analysis, and Visualization

For correlation visualization, the *ggpairs()* function was used from the GGally R package ^65^. Correlation *P*-values were corrected for multiple testing using the *corr.test()* function from the psych R package and stars (which indicate statistical significance) were hand-adjusted on the ggpairs() plot accordingly to reflect the adjusted *P*-values using BioRender. Principal component analysis (PCA) was performed using the *prcomp()* function from base R with scale set to TRUE and visualized using the *ggbiplot()* function from the ggbiplot package ^66^. Linear mixed-effects modeling was conducted using the *lmer()* function from the lme4 R package ^67^. Overall, four mixed-effects models were used to analyze NIRS-predicted macronutrient traits (protein, starch, crude fat, ash, and moisture content) as well as seed coat patterning in the RILs plus their parents (Equation 1); the same macronutrient traits in the commercial cultivars (Equation 2); the same macronutrient traits for all genotypes together (Equation 3); mineral and anti-nutrients for the subset of samples produced in Davis for which these traits were measured, and percent pigmented seed coat area in both RILs and commercial cultivars (Equation 4). In the following equations, Year signifies the planting year, Geographical Location signifies the region in which the beans were grown (coastal, inland, or intermountain), Phenotype signifies seed coat phenotypic class (and in Equation 4, also signifies the individual commercial cultivars), Phenotype:Genotype signifies RIL genotype within phenotypic class, Population Type signifies if the genotype was a commercial cultivar or a RIL, Genotype signifies the individual RIL genotype or individual commercial cultivar (within Population Type, when fitting multiple population types), Replicate signifies plot replicates in a given field trial, Environment signifies a single variable that represents the combination of the planting year and geographical location, and ε signifies the residual term. Terms that were fit as fixed effects were Genotype (in Equation 2), Population Type (in Equation 3), and Environment, Phenotype, and their interaction (in Equation 4).

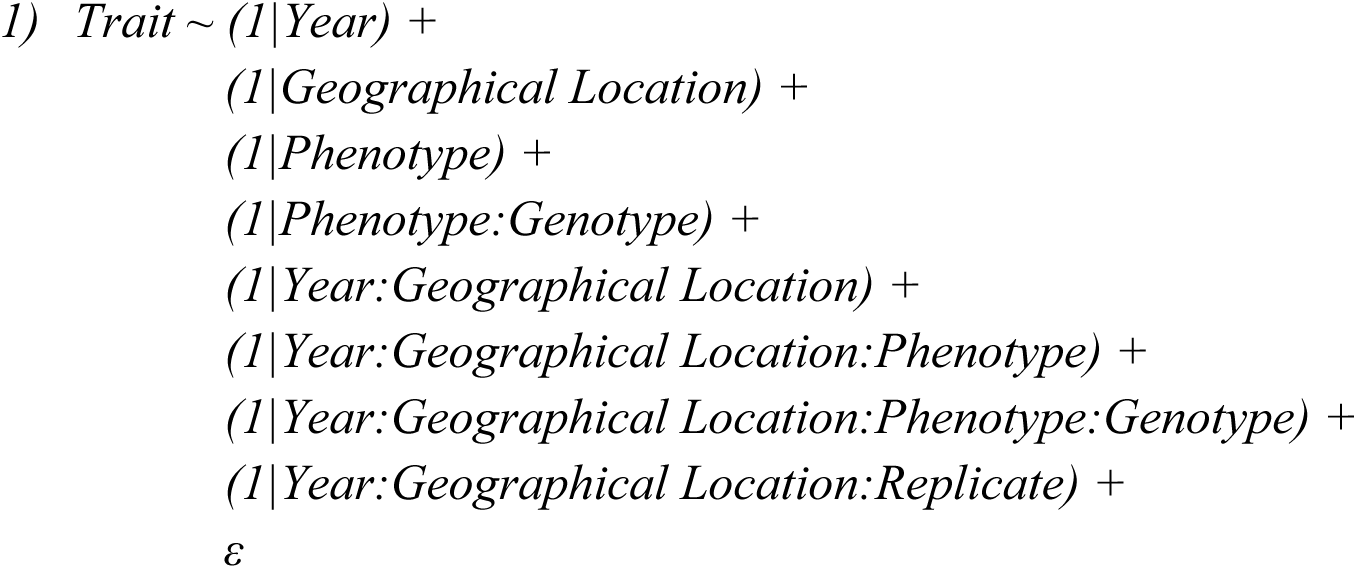

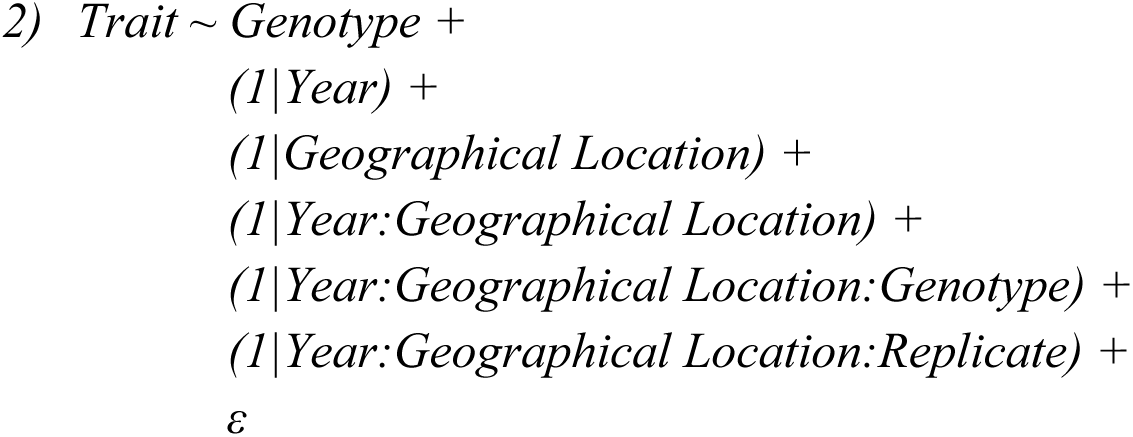

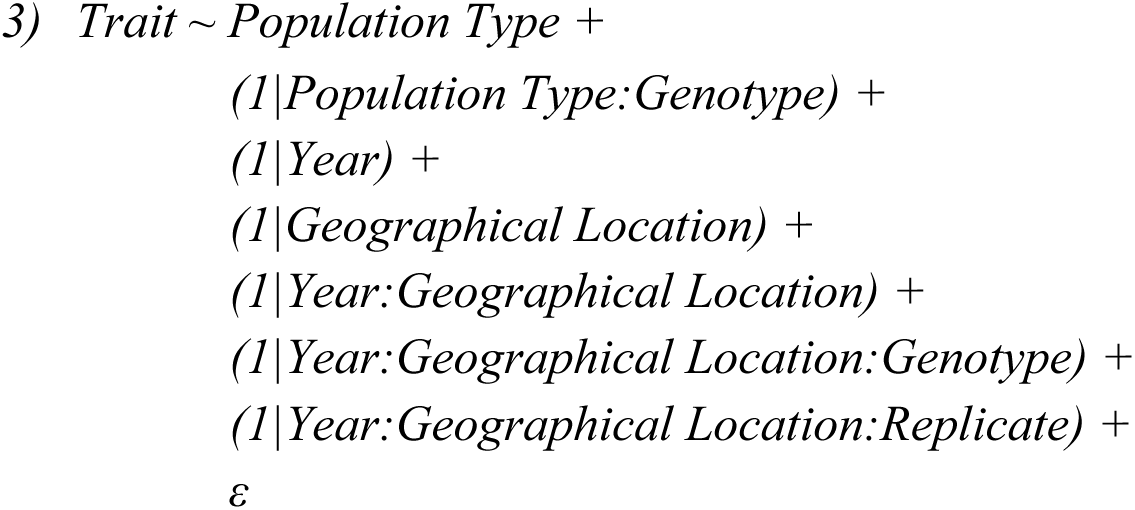

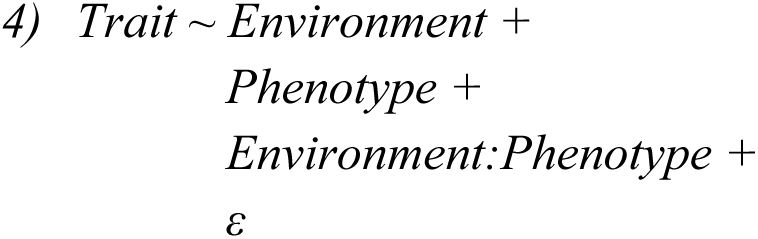

For equation 1, to estimate genotype performance of the RILs, best linear unbiased predictors (BLUPs) were extracted for each trait, with random effect estimates for the (1|Phenotype:Genotype) term obtained using the *ranef()* function from the lme4 R package. The resulting BLUPs were visualized using the *ggplot()* function from the ggplot2 R package ^64^.

For equation 2, type III ANOVA was performed using the *Anova()* function from the *car* package with use of ‘sum’ contrasts and test.statistic set to “F” ^68^. Trait means for each genotype with standard deviation were visualized using the *ggplot()* function from the ggplot2 R package^64^.

For equation 3, a linear model was fit using the *lm()* function from the lme4 R package ^67^ and was used solely to determine that the effect of Replicate was minimal prior to fitting equation 4 for percent pigmented seed coat area.

For equation 4, type II ANOVA was performed for each mineral and anti-nutrient trait in the subset of samples produced in Davis using *aov()* from base R. Group means for Phenotype were then compared using Tukey’s Honestly Significant Difference (HSD) post hoc test via the *HSD.test()* function from the agricolae package ^69^, and results were visualized using the *ggplot()* function from the ggplot2 R package ^64^. For percent pigmented seed coat area across RILs and commercial cultivars, a linear model was fit using the *lm()* function from the lme4 R package ^67^, and a post-hoc Tukey’s HSD test was conducted using the *cld*() function from the emmeans package to compare group means for the Environment:Phenotype term (calculated using *emmeans*()) at a significance level of ɑ = 0.05.

## 3 Results

### 3.1 NIRS Custom Calibration

Although performance varied across traits for the NIRS custom calibration in comparison to the pre-existing Vegetal Protein Meals (VPM) calibration, prediction accuracy improved for all traits using the custom calibration (Figure 2). For crude fat and starch, the custom calibration outperformed the VPM model, achieving a higher correlation with observed values (*r* = 0.94 vs. 0.27 and *r* = 0.77 vs. 0.48, respectively). For protein and moisture, the custom model had comparable *r* values (*r* = 1.00 vs. 0.97, and *r* = 0.90 vs. 0.86, respectively). For ash, although the custom model produced a higher prediction accuracy (*r* = 0.46 vs. 0.13), overall prediction accuracy was poor for both calibrations (*r* < 0.5). The custom calibration script was also tested for its predictive accuracy for phytate, phosphorus, iron, zinc, magnesium, calcium, potassium, and sodium content within the subset of samples produced in Davis in which these traits were measured via ICP-MS, and produced high prediction accuracies (*r* > 0.85 and *P*-value < 0.001) for iron, zinc, phosphorus, magnesium, calcium, and potassium (Figure S2).

**Figure 2.**
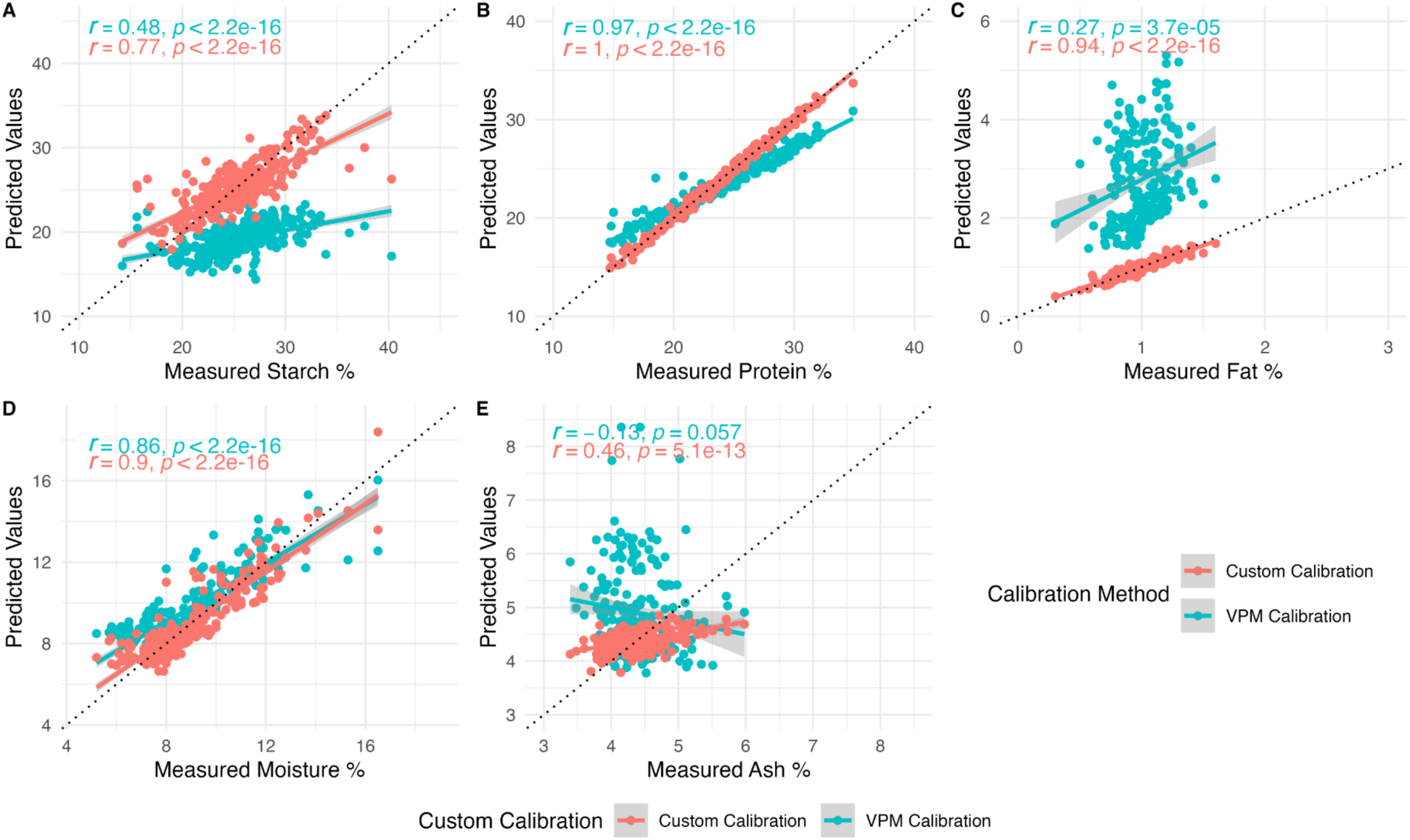
Measured (chemical reference) vs. predicted (via near-infrared spectroscopy; NIRS) values for starch, protein, fat, moisture, and ash (total mineral) percent as a percentage of seed composition. Predictions were tested using a custom NIRS calibration (in teal) or a pre-existing Vegetal Protein Meals calibration (in red). The dotted line marks the 1:1 correlation (y = x) for each trait.

### 3.2 Seed Macronutrient and Seed Coat Patterning Profiles

To explore the major axes of macronutrient trait variation across population type (i.e., commercial cultivars vs. representative RILs) and growing environments for all plots grown in 2022-2024, PCA was used (Figures 3A and 3B). Overall, the first two principal components explained 75% of total variance. In the biplot of these two PCs, ash and protein were closely aligned, indicating strong positive correlation, and were positioned opposite to starch, reflecting a strong negative correlation. Both moisture and crude fat were approximately perpendicular to each other, suggesting little to no direct correlation between them. Starch fell between moisture and crude fat, indicating an intermediate relationship to both. The RIL population displayed similar grouping in comparison to the commercial cultivars, with a slight skew towards higher ash and protein values (Figure 3A). When grouped by growing environment, the 2022 coastal growing location was the most distinct, having overall significantly higher moisture and starch and significantly lower protein content relative to any other environments (Supplemental File S1, Figures 3B and 3C). Protein content was next lowest in Intermountain 2024 and Coastal 2023 (which were not significantly different from each other), followed by Inland 2022 and 2023 and Intermountain 2023 (which were not significantly different from each other), and finally the Inland 2024 environment had significantly higher protein content relative to any other environment (Supplemental File S1). Within the inland site, protein was the only macronutrient trait that showed a significant difference across growing years; whereas all macronutrient traits showed significant differences within the intermountain site across growing years.

**Figure 3.**
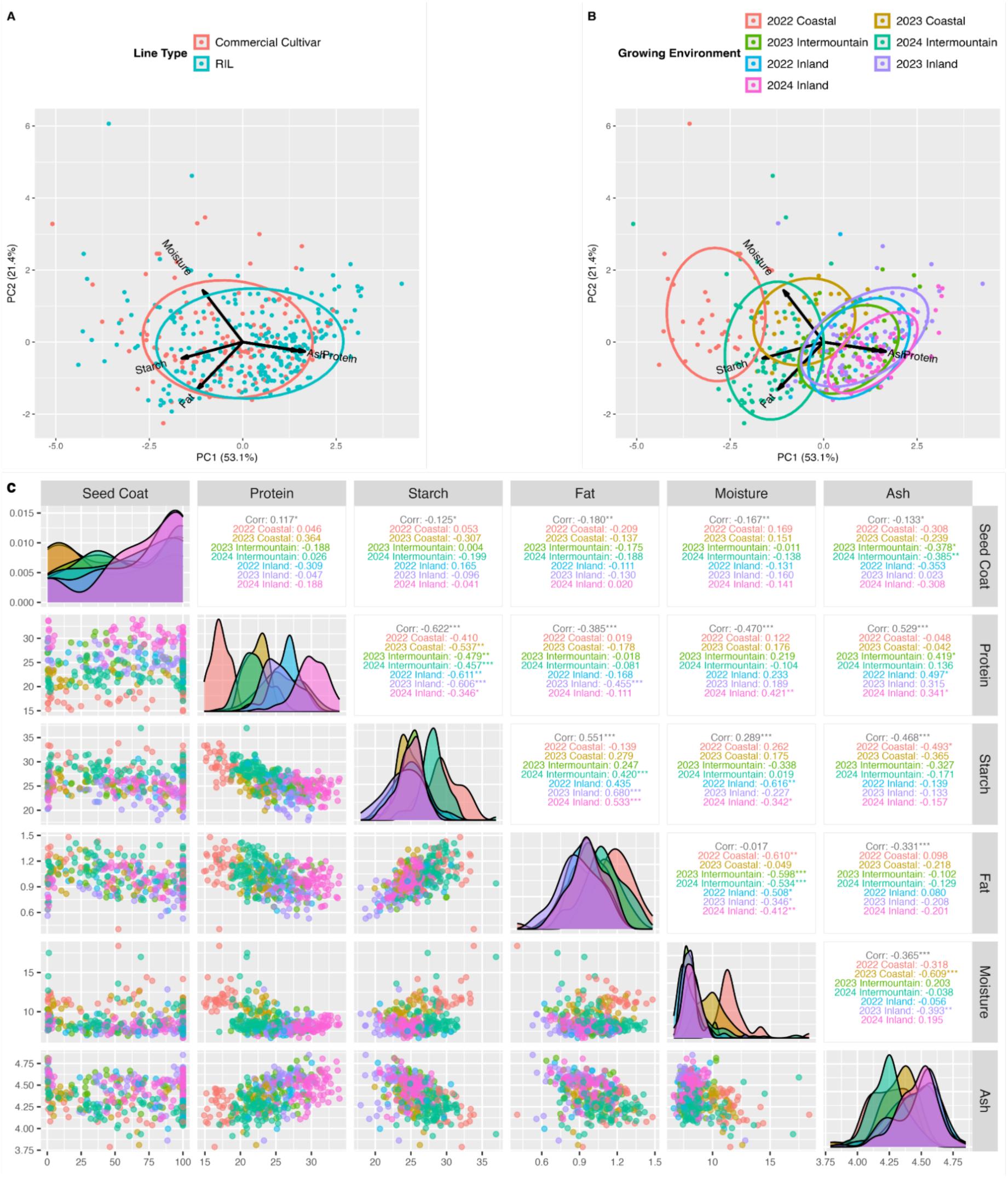
A) Biplot depicting principal components 1 vs. 2, color-coded by A) line type and B) growing environment (location by year combination). C) Trait correlation plots. The dotplots in the lower triangle and histograms on the diagonal are colored by the growing environment. The correlations in the upper triangle are the overall correlation value in black and color-coded correlation values within each growing environment (using the same color-coding as in the dotplots and histograms) with significance symbols “***”, “**”, and “*” representing adjusted *P*-values of ≤ 0.001, ≤ 0.01, and ≤ 0.05, respectively.

When examining trait distributions and pairwise trait correlations for each growing environment (Figure 3C), protein and starch displayed a significant negative correlation (averaged *r* = -0.622; *P*-value ≤ 0.001) across all growing environments. Additionally, the samples from each environment were generally organized within-environment along this negative diagonal, with some overlap in trait distributions, and with a similar organization across environments appearing for protein vs. fat. For all growing environments except for the Coastal 2023 location, moisture and fat showed a significant negative correlation (*r* = -0.346 to -0.610; *P*-value ≤ 0.05). For the Inland (2023 and 2024) and Intermountain (2024) growing locations, starch and fat were significantly positively correlated (*r* = 0.420 to 0.680; *P*-value ≤ 0.001). For the Inland (2022 and 2024) and Intermountain (2023) growing locations, ash and protein were significantly positively correlated (*r* = 0.341 to 0.497; *P*-value ≤ 0.05). For the Intermountain (2023 and 2024) growing locations, ash and percent pigmented seed coat area were significantly negatively correlated (*r* = -0.378 to -0.385; *P*-value ≤ 0.05). For the Coastal (2022) growing location, ash and starch were significantly negatively correlated (*r* = -0.493; *P*-value ≤ 0.05).

Coastal 2022 yield was significantly correlated with protein content in the mixed population (*r* = -0.45 and *P*-value < 0.01); however, when parsed by population type, this relationship was only significant in commercial cultivars (*r* = -0.88 and *P*-value < 0.01) and not in RILs (*P*-value > 0.05) (Figure S3). Inland 2024 yield was not significantly correlated with protein content (*P*-value *>* 0.05; Figure S3).

For the RILs and the five commercial cultivars with partial seed coat pigmentation, strong variation in percent pigmented seed coat area across growing environments was observed (Figure 4). For commercial cultivars with partial seed coat patterning in phenotypic class 3 (Anasazi, UCSW Red, UCSW Gold, and Orca), the least pigmented seed coat area was observed on seed grown coastally, with intermediate values observed in the intermountain location, and the highest pigmented seed coat area was observed on seed grown inland. For RILs of phenotypic class 3, this trend in pigmented seed coat area across locations was also observed. Samples in phenotypic class 4 (including RILs and Black Nightfall), which exhibit a speckled seed coat pattern, showed an opposite trend to phenotypic class 3 across locations, with the lowest percent pigmented seed coat area observed in the inland 2022 and 2023 growing environments. RILs in phenotypic class 4 also had particularly high percent pigmented seed coat area (with narrow variability) in the 2023 coastal location, which was significantly higher than in five of the other six environments for this phenotypic class. Phenotypic class 5 followed a trend more similar to phenotypic class 4 than to 3, though with fewer significant pairwise combinations across environments than those observed within phenotypic class 4. For RILs in phenotypic class 2, the coastal 2022 growing environment produced significantly less percent pigmented seed coat area than inland 2023 and 2024, though no other significant differences were observed. As expected, the RILs in phenotypic classes 1 and 6 displayed the least amount of variation (and no significant differences) across growing environments, as they did not show partial seed coat pigmentation.

**Figure 4.**
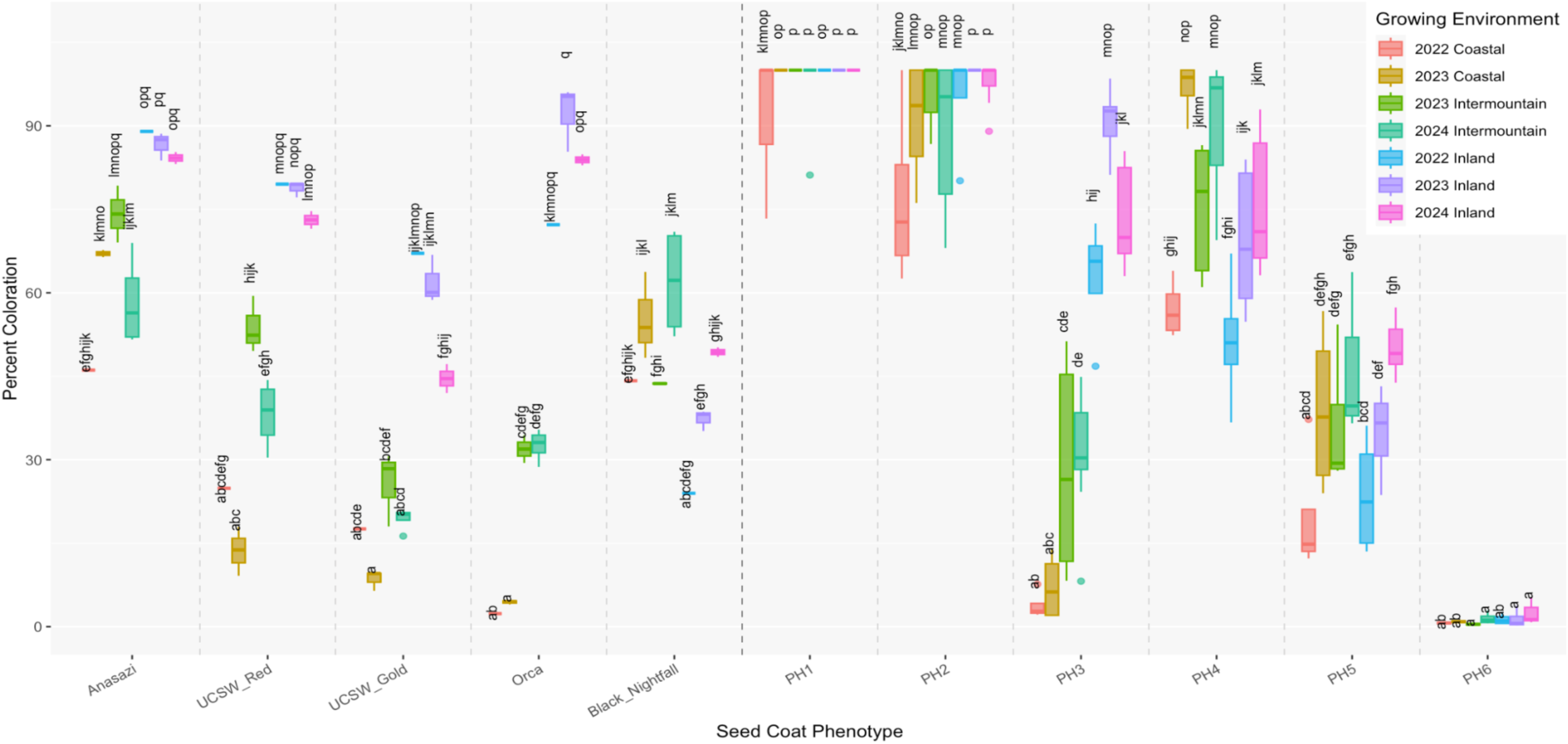
Boxplots of the percent pigmented seed coat area for commercial cultivar genotypes with partial patterning as well as all phenotypic classes of the RIL population, sorted by year and location grown. Each genotype is colored by growing location and year. The dashed line separates the commercial cultivars (left) from the RILs (right). Post-hoc analysis was done for the commercial cultivars and the RILs as two separate groups; samples sharing no letters were significantly different from each other (adjusted *P*-value < 0.05).

When evaluating macronutrient profiles of the commercial cultivars, the effects of genotype, growing location, and their interaction were significant (all *P*-values < 0.01) for all traits tested (protein, starch, fat, moisture, and ash), with the exception that the interaction term was not significant for starch (Supplemental File S1). The commercial cultivars generally retained their rank order for protein across environments, with Black Nightfall and Orca consistently showing high levels and Viva consistently showing low levels (Figure 5A). Additionally, for protein, year and its interaction with each of genotype and location had a significant effect (all *P*-values < 0.01) with a range of genotypic means of up to 10.29% on the original trait scale (i.e., 19.67% protein for Viva to 29.96% protein for Black Nightfall in Intermountain 2023). For starch, variation was low (with a range of mean values across genotypes of 3.22 to 7.39%) within a given growing environment, except for Inland 2022 (which had a 9.97% range), and the effect of year was not significant (Figure 5B and Supplemental File S1). Among the commercial cultivars, Anasazi consistently had low values for fat (particularly in Inland 2022 and 2023; 0.62 and 0.64%, respectively, with all other genotypes ≥ 0.80%), low values for ash (particularly in Coastal 2022; 3.79%, with all other genotypes ≥ 4.08%), and high values for moisture (11.01 and 10.51% in Inland 2022 and 2023, with all other genotypes < 8.57% in both years; and 13.46% in Intermountain 2024, with all other genotypes < 9.31%) (Figures 5C, 5D, and 5E).

**Figure 5.**
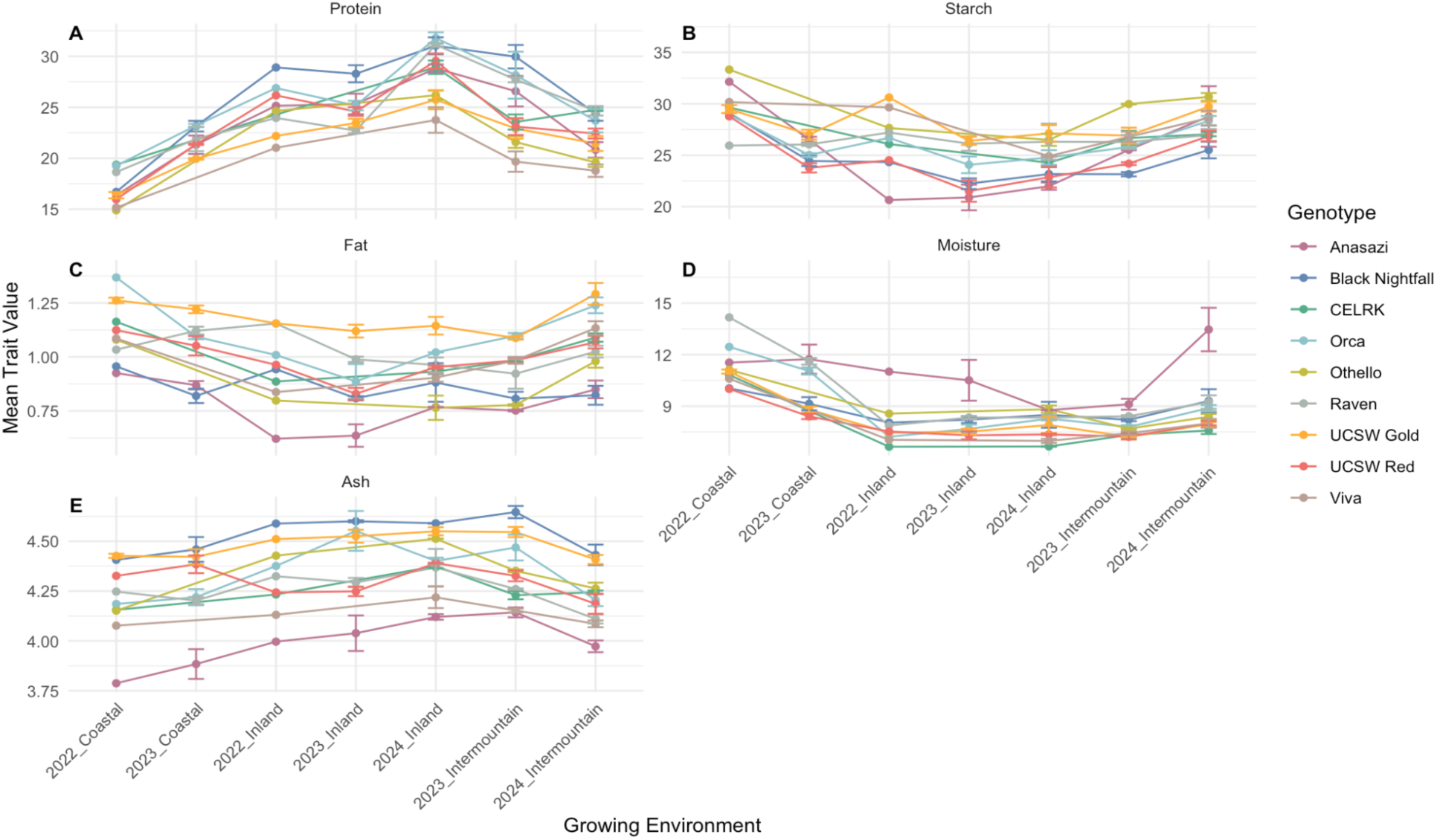
Commercial cultivar trait values for (A) protein, (B) starch, (C) fat, (D) moisture, and (E) ash (total mineral) percent, as a percentage of seed composition, in each growing environment. Points are mean values for each trait with vertical lines depicting standard deviation across plots within a growing environment (except in 2022, when samples were pooled within location) and connecting lines depicting the same genotype across environments.

Evaluating the RIL BLUPs grouped by phenotypic class showed high variability in genetic potential for percent pigmented seed coat area as expected, with a particularly wide range (-23.3 to 16.5) for phenotypic class 4 and narrow ranges for phenotypic classes 1 and 6 (-0.1 to 1.4 and -2.0 to -0.3, respectively) (Supplemental File S1, Figure 6). Notably, for protein, phenotypic class 1 had a narrow BLUP range centered around zero while all other phenotypic classes were more wideranging (i.e., BLUPs for RILs in phenotypic class 3 ranged from -1.4 to 1.9).

**Figure 6.**
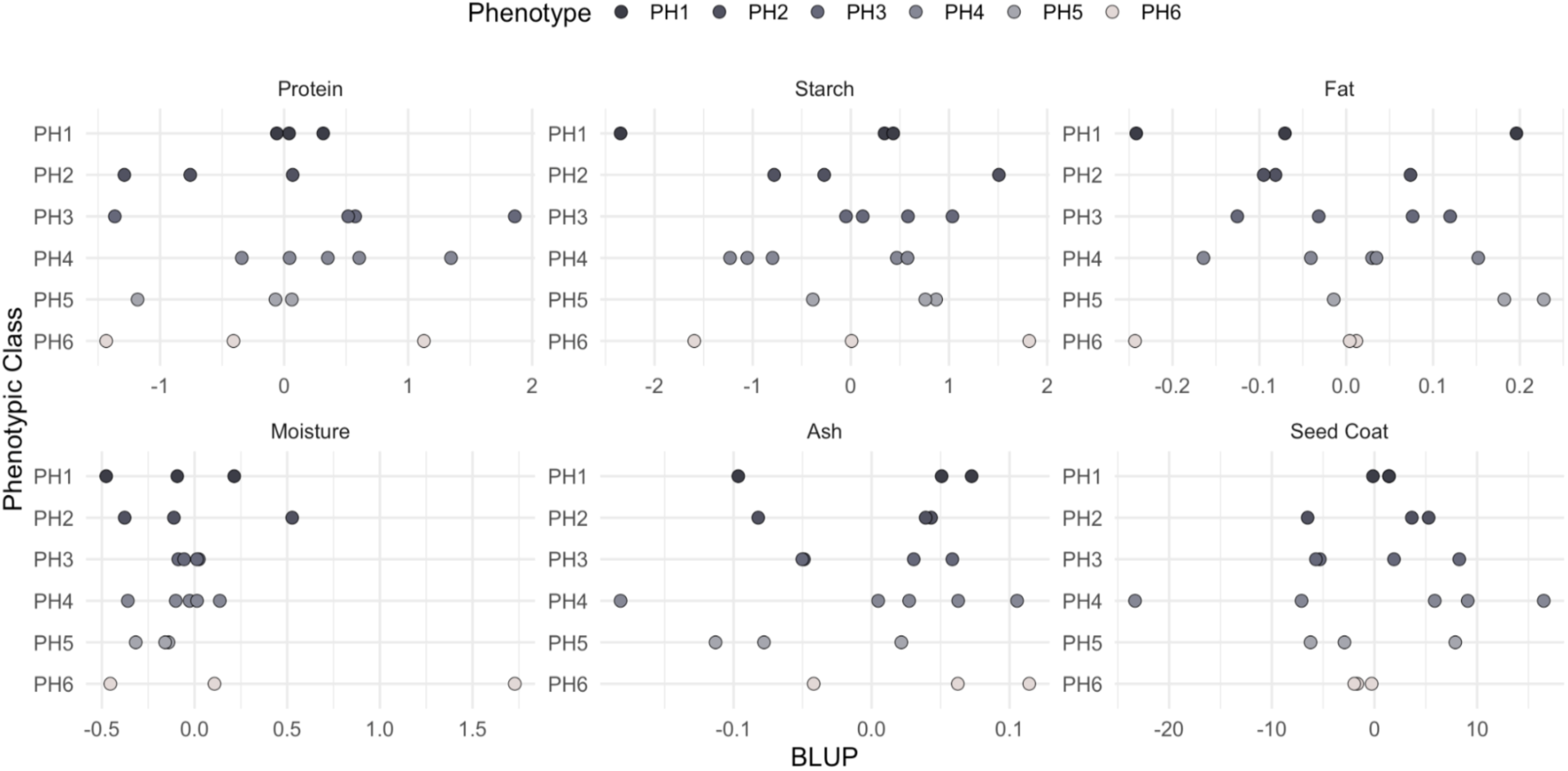
RIL and parent cultivar (Black Nightfall and Orca) best linear unbiased predictors for protein, starch, fat, moisture, ash (total mineral), and percent pigmented seed coat area. Dots represent individual genotypes and are colored by seed coat phenotypic class.

Genotype OxB-094 from phenotypic class 3 displayed a particularly high BLUP value (1.9) for protein (Supplemental File S1). Genotype BxO-070 from phenotypic class 1 had the lowest BLUP value for starch (-2.3). Additionally, starch BLUPs in phenotypic class 3 were shifted toward positive values (ranging from -0.05 to 1.03). Both ash and fat BLUPs showed little class-level structure, with most phenotypic classes exhibiting a mixture of positive and negative values and no consistent directional trends. Phenotypic class 5 was an exception, displaying a modest shift toward higher fat BLUPs. Moisture content exhibited clear differences among phenotypic classes. Phenotypic class 3 was tightly clustered around zero, reflecting minimal within-class variation. The most striking pattern for moisture was observed in phenotypic class 6, which included one genotype (OxB-179) with an extreme positive BLUP (1.7), exceeding the range observed in all other classes by three fold.

### 3.3. Soil Analysis and Seed Micronutrient and Anti-nutrient Profiles

Soil samples taken from the Inland growing location in 2022, 2033, and 2024 were analyzed for pH and mineral nutrients (Table 1). Although slight differences were observed, there were no significant differences (all *P*-values > 0.05) in these traits between the sampling years.

**Table 1.** Soil analysis results for Davis 2022, 2023, and 2024 for pH and mineral nutrient concentrations in µg per g for iron (Fe), zinc (Zn), calcium (Ca), magnesium (Mg), phosphorus (P), potassium (K), and sodium (Na). Averages and standard deviations for samples from three replicate field plots from each year.

| Year | Soil pH | Fe (µg/g) | Zn (µg/g) | Ca (µg/g) | Mg (µg/g) | P (µg/g) | K (µg/g) | Na (µg/g) |
| --- | --- | --- | --- | --- | --- | --- | --- | --- |
| 2022 | 7.00 ± 0.32 | 33079.37 ± 3657.44 | 76.64 ± 9.08 | 3641.99 ± 387.67 | 18610.00 ± 2481.89 | 446.33 ± 50.62 | 3357.63 ± 316.14 | 157.45 ± 9.81 |
| 2023 | 7.08 ± 0.08 | 34488.63 ± 1033.82 | 76.31 ± 2.95 | 3819.15 ± 118.54 | 19933.18 ± 1090.97 | 442.74 ± 22.66 | 3283.25 ± 468.97 | 153.49 ± 17.98 |
| 2024 | 6.78 ± 0.09 | 35628.75 ± 184.50 | 81.48 ± 1.55 | 3897.62 ± 102.07 | 18525.26 ± 252.38 | 463.76 ± 7.22 | 3820.81 ± 211.66 | 134.55 ± 5.27 |

Seed phytate content and mineral nutrient concentrations were only measured in the Inland environment (in 2022, 2023, and 2024) due to the availability of soil data from those three trials. For phytate, phosphorus, zinc, potassium, and sodium, the growing year was significant (all *P*-values < 0.05). Compared to any other year, phytate and phosphorus were significantly higher in 2023; zinc was significantly lower in 2022; and potassium was significantly lower in 2024 (Supplemental File S1). Sodium was significantly higher in 2024 than 2022. For phosphorus, iron, zinc, magnesium, and calcium, phenotypic class (for RILs) or commercial cultivar was significant (all *P*-values < 0.05). For phytate versus phosphorus, the rank order of genotypes (commercial cultivars and the RILs grouped by phenotypic class) was largely preserved (Figures 7A and 7B). Additionally, significant differences between genotypes were detected for iron, zinc, and magnesium (Figures 7C, 7D, and 7E). Black Nightfall ranked in the top four for iron, zinc, phosphorus, and magnesium but was second-to-last for calcium. Additionally, the phenotypic class 4 RILs (the same class as Black Nightfall) performed within the top six for iron, zinc, and magnesium. Weak performers for iron, zinc, phosphorus, and magnesium included Viva, UC Southwest Gold, Anasazi, and Raven (with the exception of magnesium levels in UC Southwest Gold being on the higher end and Raven being mid-range for magnesium). For phytate, calcium, potassium, and sodium there were no significant differences between the genotypes, and for the latter three traits, rank order was loosely preserved for only a few genotypes (i.e., CELRK was generally on the lower end while Orca was generally on the higher end) (Figures 7F, 7G, and 7H).

**Figure 7.**
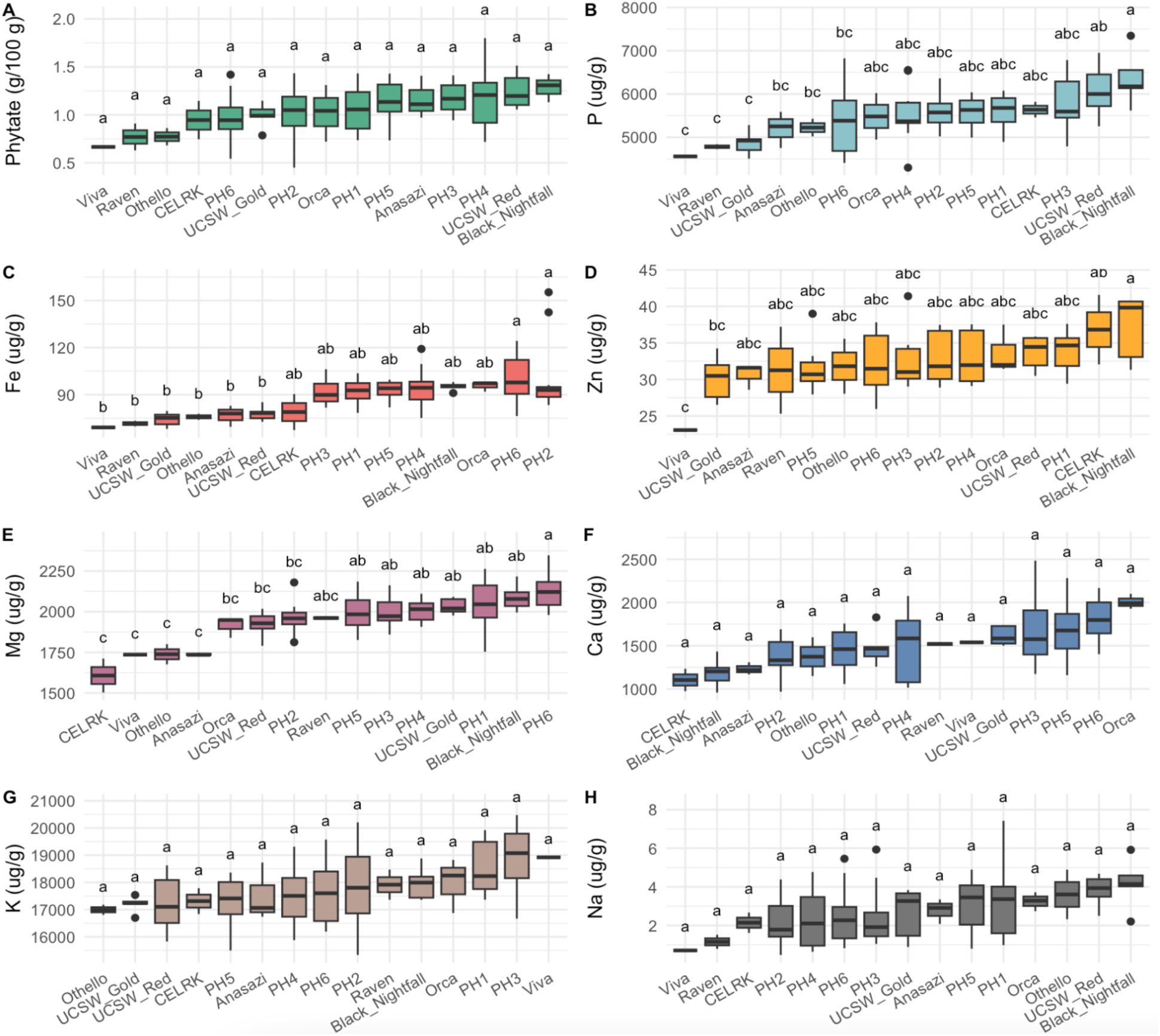
For seed samples from field plots in Davis from 2022, 2023, and 2024: (A) phytate content in g per 100 g and mineral nutrient concentrations in µg per g for (B) phosphorus (P), (C) iron (Fe), D) zinc (Zn), E) magnesium (Mg), F) calcium (Ca), G) potassium (K), and H) sodium (Na), sorted in ascending order of trait value within each panel and labeled via seed coat phenotypic class (for RILs) or commercial cultivar name.

To examine how all measured traits relate to one another and to assess broader multivariate patterns across years within the Inland growing location (the only location for which all traits were measured), a combined correlation matrix and two PCAs were generated (Figure 8). To visualize the pairwise correlations among all seed traits for the samples grown in the Inland location across all growing years, traits were clustered into three main groups: seed coat, macronutrients, phytate, and mineral nutrients (Figure 8A). Macronutrients which displayed significant positive correlations included protein vs. moisture and starch vs. fat. A significant negative correlation was observed between starch and protein as expected, and between moisture and both starch and fat. Additionally, there was a significant negative correlation between starch and phytate. Within mineral nutrients, many significant positive correlations were observed, with zinc and phosphorus having the strongest association (*r* = 0.54; *P*-value < 0.001). Potassium and calcium were the only mineral nutrients that were significantly negatively correlated with each other. Cross-category correlations (e.g., between macronutrients and specific minerals) resulted in a few notable associations, indicating that some minerals covaried with particular macronutrients. A positive correlation was observed between protein and zinc, and percent seed coat coloration was negatively correlated with each of magnesium and calcium. As expected, a significant strong positive correlation was observed between phytate and phosphorus.

**Figure 8.**
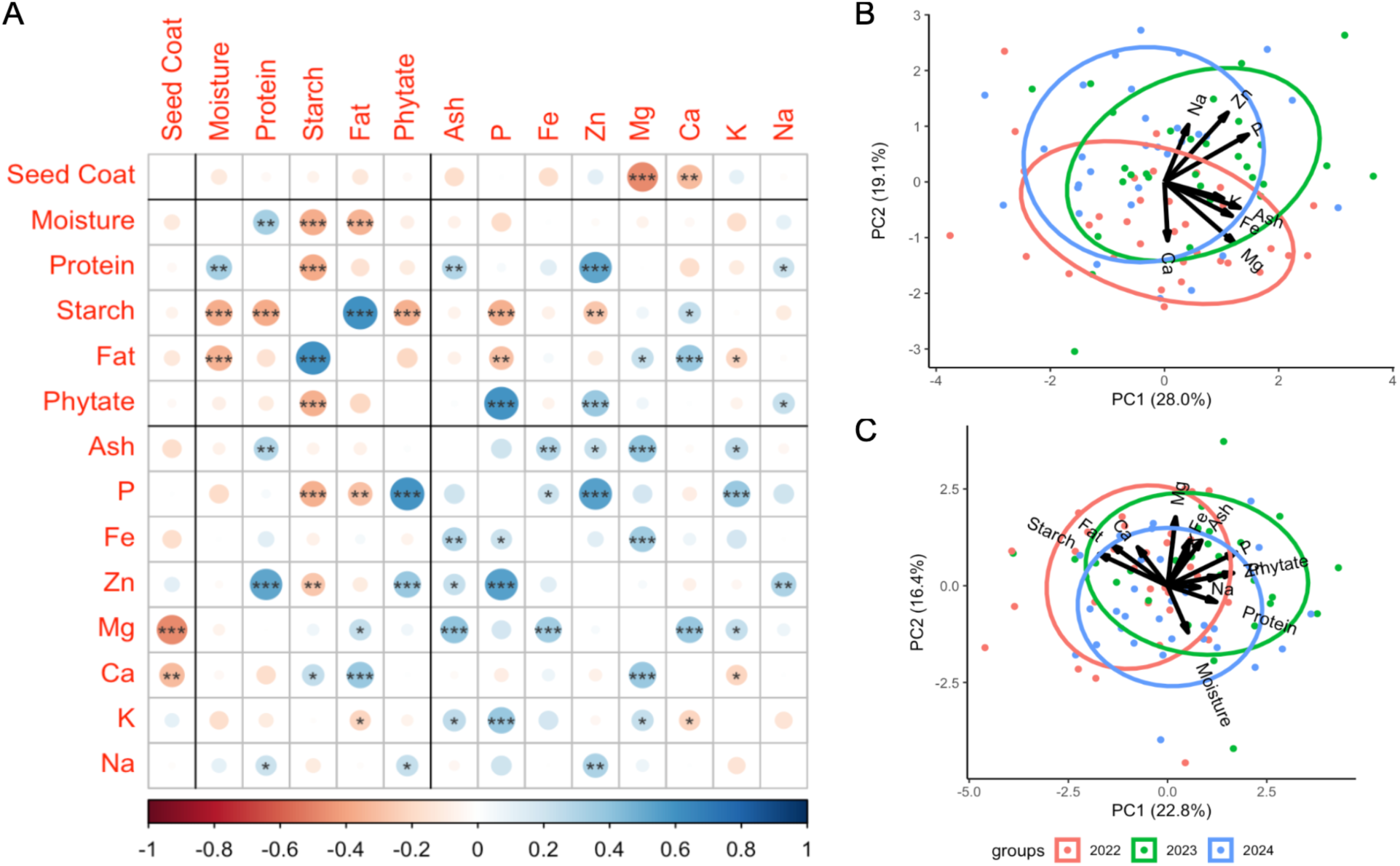
For seed samples from field plots in Davis from 2022, 2023, and 2024, which were selected for phytate and mineral nutrient analysis: A) trait correlations with all data including seed coat, macronutrients, phytate, and mineral nutrients (phosphorus (P), iron (Fe), zinc (Zn), magnesium (Mg), calcium (Ca), potassium (K), and sodium (Na)) with the red-blue color scale representing *r* value and significance symbols “***”, “**”, and “*” representing *P*-values of ≤ 0.001, ≤ 0.01, and ≤ 0.05, respectively. B) PCA for mineral nutrient samples separated by year. C) PCA for samples with all data separated by year.

The PCA using only mineral nutrients and ash content for the samples grown in the Inland location shows moderate multivariate structure, with the first two principal components (PC1 and PC2) explaining 47.1% of the total variance (Figure 8B). The growing years formed partially overlapping but distinguishable clusters (particularly samples from 2022), suggesting environmental conditions influenced mineral nutrient profiles in a consistent manner across samples. The incorporation of all measured macronutrients and mineral nutrients for the samples grown in the Inland location yielded a similar multivariate pattern (Figure 8C). However, the combined nutrient dataset decreased the overall variation captured by PC1 and PC2 (39.2%), and the growing years formed less discernible clusters. Variance component analysis showed a minor effect of growing year for most traits, with the exception of protein (50.79%) and zinc (44.20%) (Supplemental File S1). However, even with this by-year variation, the inland growing location generally produced the highest protein content (Figure 3C).

## 4 Discussion

### 4.1 Nutritional Quality Traits and Genotypes for Breeding Enhancement

Among the commercial cultivars evaluated under this GxE framework, Black Nightfall and Orca displayed stable high protein content (consistently ranking in the top four among commercial cultivars) across environments. These characteristics suggest that Black Nightfall and Orca could be candidate parents for use in breeding programs targeting protein improvement. Additionally, of the RILs produced by crossing Black Nightfall and Orca, genotype OxB-094 from phenotypic class 3 displayed a particularly high protein BLUP value (1.9), indicating that this genotype could be a parental candidate for protein enhancement in breeding programs and for dissecting the genetic architecture underlying this trait (Supplemental File S1). Viva consistently displayed low protein content (lowest-ranking in all but one environment; though there could be a protein-yield tradeoff as Viva exhibited high yield in the Coastal 2022 environment, and its high yield index has been previously noted ^52^). Total protein content is not necessarily indicative of the total content of quality protein. Breeding strategies that target specific improvements in protein quality (e.g., digestibility ^20,70,71^ and amino acid composition) may help mitigate tradeoffs between total protein and yield ^72–75^.

Although phenotypic class 1 had a narrow BLUP range for protein, genotype BxO-070 from this class had the lowest BLUP value for starch (-2.3). This pattern suggests that starch and protein were not varying in parallel within this class, and that individual genotypes may have differed in their allocation of carbon and nitrogen into storage compounds ^76^. Notably, genotype OxB-179 from phenotypic class 6 had an extreme positive BLUP (1.7) for moisture, exceeding the range observed in all other classes by three-fold. This extreme value highlights substantial genetic variability for moisture and identifies a potential target for further investigation given the agronomic relevance of seed moisture to harvest timing, post-harvest handling, and storage stability ^77^. In contrast, the BLUP results for fat and ash (total mineral) content suggest that these traits were not strongly structured by RIL seed coat phenotypic class. Importantly, the RILs examined herein were a subsample from a larger population. Examination of more genotypes within and across phenotypic classes may help in assessing systematic trends and parsing more subtle differences in macronutrient profiles.

### 4.2 Mineral Composition, Seed Coat Traits, and Environment

In comparison to previous literature for common bean, the genotypes tested herein displayed comparable mineral nutrient levels ^78,79^. Correlation and PCA analysis on mineral nutrients were indicative of certain mineral nutrients co-varying more closely (particularly for zinc and phosphorus) than others (Figure 8). Calcium and magnesium were of particular interest as they have been previously associated with preferential partitioning to the seed coat, where magnesium and especially calcium are enriched relative to their distribution in whole-seed dry matter ^80^, and with cooking quality ^81^. While Hacisalihoglu et al. found that common bean with black seed coats displayed high levels of mineral content (including magnesium), our study found a significant negative correlation between percent of pigmentation in the seed coat and each of calcium and magnesium concentrations (Figure 8A)^82^. This contrast may reflect differences in experimental design, as their study did not include partially pigmented seed types but did compare the black beans against a broader range of seed coat colors. Previous studies have also shown substantial environmental effects on seed iron and zinc concentrations as well as seed coat coloration/patterning ^16,17^. For example, the RILs and commercial cultivars with partial seed coat pigmentation significantly differed in percent pigmented seed coat area across growing environments. This was as expected and in correspondence with previous research which found that higher temperatures resulted in higher pigmented seed coat area (Figure S1)^16^. Although mineral concentrations were only explored for one growing location herein across three years, extending future work to include calcium and magnesium assessment in multi-location trials would further our collective understanding of the accumulation of these seed minerals of relevance for human health in a GxE context.

### 4.3 Monitoring and Management of Anti-Nutrients

This study highlights challenges in the assessment of anti-nutritional factors such as phytate. As expected, phytate showed a strong positive correlation with phosphorus, reflecting its role as the primary phosphorus storage compound in seeds. Although no significant differences by phenotypic class were observed for seed phytate content when measured via colorimetric assay, significant differences in seed phosphorus levels were observed. The significant negative correlation observed for phytate (and phosphorus) vs. starch is in agreement with Wang et al., who reported a significant correlation of -0.48 between phytate and starch in 20 cultivars and breeding lines that were evaluated in four field environments in Manitoba, Canada ^83^. These findings highlight the need to consider both nutritional and anti-nutritional content and potential relationships between and among them when defining breeding targets.

Additional anti-nutritional and bioactive compounds, such as phenolics, also warrant further investigation, particularly within multi-environment trials that capture their environmental sensitivity. In common bean, phenolic compounds contribute to seed coat patterning ^84,85^ and nutritional quality, including interactions with both starch ^86–88^ and proteins^89^. Thus, integrating phenolic profiling into GxE frameworks will be critical for understanding how environmental variation shapes both nutritional quality and health outcomes ^90–94^.

### 4.4 Genotype by Environment

In the Inland location, the same amount of nitrogen was applied in 2022 and 2024, and more nitrogen was applied in 2023 due to weed pressure. The 2022 and 2023 trials were in the same field block, which has a Reiff very fine sandy loam soil type, and the 2024 trial was in a different field block, which has a Yolo silt loam soil type ^95,96^. For RILs, the 2024 Inland environment had significantly higher protein content compared to any other environment tested in this study; however, within the Inland location for all other macronutrient traits (starch, fat, ash, and moisture), no significant differences were observed. Additionally, for RILs, the Intermountain 2024 had significantly lower protein than 2023, despite similar N application (110 and 100 lb per acre, respectively). In the mixed population, significantly higher seed phytate and phosphorus levels were observed in Inland 2023, which also had a higher fertilizer (nitrogen-phosphorus-potassium) rate, than any other year. However, soil levels of nitrogen, phosphorus, and potassium were not significantly different across years, and seed potassium was not significantly different between 2022 and 2023 and was significantly lower in 2024. These trends in grain mineral nutrient profiles could have been due to multiple factors that could influence nutrient uptake including applied nitrogen, residual nitrogen in the field, weed pressure, and soil type, and/or could reflect differences in nutrient transport to the seed ^97,98^. Examination of mineral nutrient concentrations in multiple tissues of the plant through the growing season would help in further resolving the underlying mechanisms with relevance to both breeding and agronomy.

Further, examination of more representatives within each of the major market classes of dry beans and full biparental and multi-parental populations from multi-environment trials for macronutrient, anti-nutrient, and mineral nutrient traits would be informative in assessing GxE effects, genetic underpinnings, and favorable variation to be leveraged in breeding. Lastly, seed zinc concentrations in Inland 2022 were significantly lower than in any other growing year. This may reflect environmental constraints on zinc uptake or remobilization at grain filling ^97,98^.

Examination of micronutrient-specific GxE responses and their physiological drivers would be informative when targeting stable seed mineral density in breeding programs.

### 4.5 Custom NIRS Calibrations and Expanded Application

Major limitations on routinely profiling nutritional composition alongside agronomic traits in multi-environment field trials are the time and cost involved in accurately measuring macro- and micronutrient composition. Near-infrared spectroscopy can be a powerful tool for prediction of macronutrient composition, but it is imperative to use a calibration that is representative of the sample set being predicted. Many pre-existing calibrations exist but are limited to a narrow range of crop commodities. For example, the existing VPM calibration used in this study included legumes but was primarily trained on oilseeds and meals derived from them, and did not adequately represent the nutrient composition of common bean, resulting in predictions that were particularly suboptimal for starch (VPM *r* = 0.48), fat (VPM *r* = 0.27), and ash (VPM *r* = -0.13) (Figure 2). By developing a custom calibration, prediction accuracy was improved for all macronutrient traits but particularly for starch and crude fat. While an intermediate approach of augmenting the existing calibration with a subset of study-specific samples could have also conferred some improvement, such an approach was not feasible due to software constraints.

Overall, with chemical analyses for 220 samples, prediction accuracies of *r* = 1.00, 0.77, 0.94, 0.90, and 0.46 for protein, starch, fat, moisture, and ash respectively were obtained for the full sample set of 394 using the custom calibration. Additionally, for the traits with final prediction accuracies below 0.90 (starch and ash), small differences between samples may have been hidden by larger prediction error; further improvement of these prediction accuracies via updated modeling approaches (e.g., machine learning algorithms ^99^) or higher-resolution chemical analysis (e.g. for individual carbohydrates ^100,101^) may unveil more subtle differences between the samples.

Additionally, recent studies have explored the use of NIRS for indirect prediction of mineral nutrients ^102–104^. For common bean samples, Garcia et al. used NIRS to detect iron and zinc in whole seeds, and Plans et al. used NIRS to detect calcium and magnesium in whole seeds and ground seed coats ^105,106^. Although those studies found that NIRS-based predictions for iron and zinc were less efficient than for nitrogen, like nitrogen, these minerals interact with organic compounds, enabling their indirect estimation ^106^. High accuracies of NIRS-based predictions for iron, zinc, phosphorus, magnesium, calcium, and potassium (based on chemical reference values from ICP-MS) were also observed in this study, reinforcing the utility of NIRS for rapid mineral profiling (Figure S2). However, NIRS prediction accuracy was poor for phytate, which was quantified using colorimetric assays (Figure S2). Collectively, these findings suggest that NIRS could serve as a valuable tool for mineral assessment in breeding programs, particularly when coupled with robust calibrations adapted for the sample set under study. However, given the reliance on chemical bonds for prediction, it would be important to ensure that the predictable mineral concentrations also tend to be bioaccessible/bioavailable during digestion, which also depends on seed coat coloration and patterning due to phenolic-mineral nutrient interactions. Overall, use of NIRS could be helpful in scaling the routine and cost-effective analysis of macronutrients and potentially also mineral nutrients out to the multi-environment field and population scales.

### 4.6 Conclusions

Overall, this study demonstrates substantial genetic and environmental influences on seed coat patterning, macronutrient and phytate content, and mineral concentrations in common bean, highlighting both opportunities and challenges for breeding pipelines and the food supply/demand streams. Identification of high-protein cultivars and RIL genotypes provides a foundation for improving nutritional quality. At the same time, complex relationships among protein, starch, minerals, and phytate underscore the need for integrated selection strategies for overall optimized nutritional value. Additionally, seed coat patterning (a visual characteristic that majorly defines market class and influences consumer preference) had significant correlations with both calcium and magnesium. Advances in high-throughput assays, particularly the development of robust NIRS calibrations, offer practical pathways for scaling these efforts in breeding programs. To fully realize the contributions of common bean to agricultural and food systems, nutritional composition must be routinely measured in samples from multi-environment trials. This effort should extend to a broader suite of nutritional and anti-nutritional traits and incorporate integrated, multi-trait selection strategies that account for correlations among macronutrients, minerals, and anti-nutrients, with explicit evaluation of GxExManagement effects.

## Acknowledgements

We gratefully acknowledge Jonathan Berlingeri, Mary-Francis LaPorte, Jaclyn Adaskaveg, and Sassoum Lo for the collaborative work put into developing the custom NIRS calibration script. We also gratefully acknowledge the field station teams at UC Davis, the UC Santa Cruz Center for Agroecology Farm, and the UC ANR Intermountain Research & Extension Center. We also gratefully acknowledge Arnaldo Rios-Cruz for assisting in the collation of initial seed stocks.

## Author contributions: CRediT

**Tayah M. Bolt** - Conceptualization, Data curation, Formal analysis, Investigation, Methodology, Project administration, Resources, Software, Supervision, Validation, Visualization, Writing – original draft, Writing – review and editing

**Austin Cole** - Investigation, Resources, Writing – review and editing

**Rajdeep Bains** - Investigation, Writing – review and editing

**Li Tian** - Funding acquisition, Writing – review and editing

**Travis A. Parker** - Project Administration, Funding acquisition, Resources, Writing – review and editing

**Paul Gepts** - Funding acquisition, Resources, Writing – review and editing

**Antonia Palkovic** - Project Administration, Resources, Writing – review and editing

**Gail M. Bornhorst** - Funding acquisition, Resources, Writing – review and editing

**Christine H. Diepenbrock** - Conceptualization, Formal analysis, Funding acquisition, Methodology, Project administration, Resources, Software, Supervision, Validation, Visualization, Writing – original draft, Writing – review and editing

---

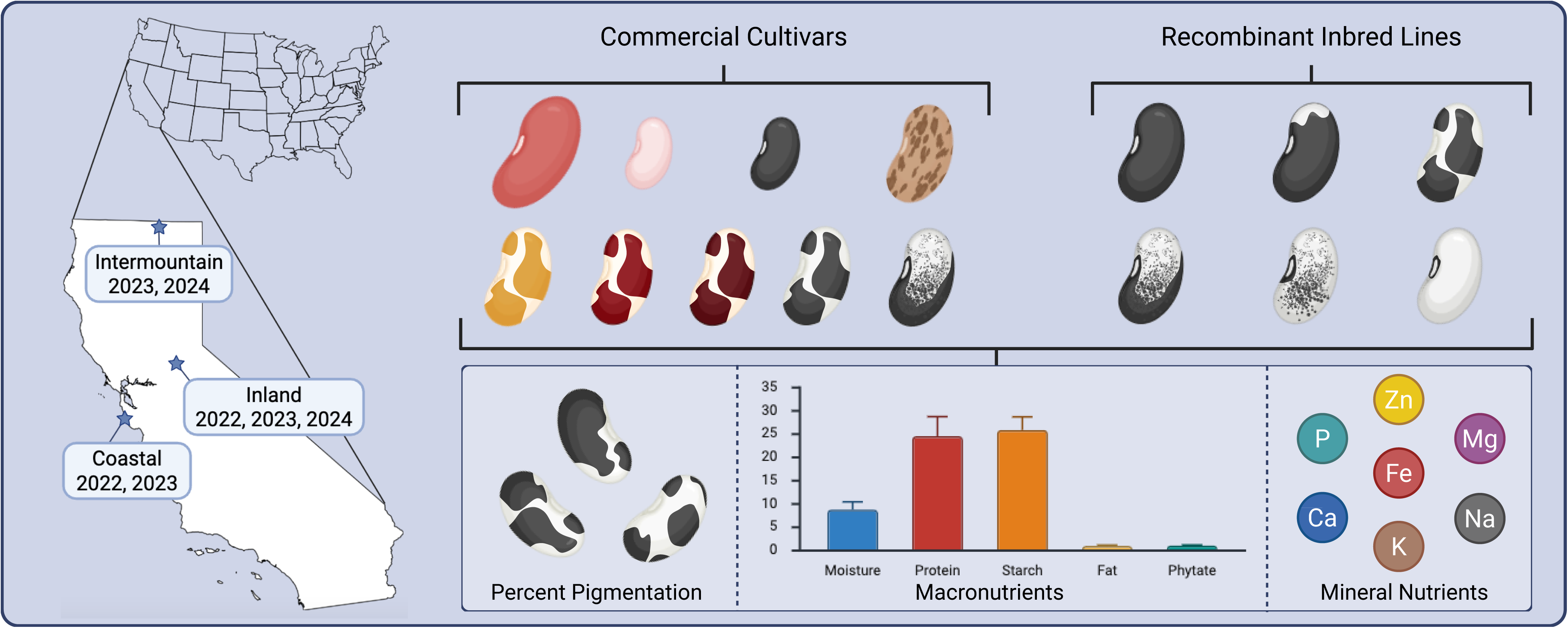

