## Supplemental Figures and Tables for "Genotypic and environmental effects on seed coat patterning and nutritional composition in common bean (*Phaseolus vulgaris* L.)"

Table S1. Common bean genotypes planted for small-plot trials across California in 2022, 2023, and 2024.

<table>
<tr><th>Year</th><th>Growing Locations</th><th>Planting Date</th><th>Harvest Date</th><th>Plots per Genotype</th><th>Genotypes Planted</th></tr>
<tr><td rowspan="2">2022</td><td>Davis, CA</td><td>May 25th</td><td>September 6-7th</td><td>3</td><td rowspan="2">Commercial cultivars: Anasazi, Black Nightfall, CELRK, Orca, Othello, Raven, UCSW Gold, UCSWRed, Viva<br>Phenotypic Class 1: BXO-030, OXB-098, OXB-100, OXB-178<br>Phenotypic Class 2: BXO-119, BXO-144, BXO-158, OXB-086<br>Phenotypic Class 3: BXO-052, BXO-139, BXO-0145, OXB-094<br>Phenotypic Class 4: BXO-040, BXO-049, BXO-055, BXO-145<br>Phenotypic Class 5: BXO-100, BXO-118, BXO-159, OXB-117<br>Phenotypic Class 6: BXO-013, BXO-075, OXB-139, OXB-179<br>San Gregorio Border: UCSW Gold</td></tr>
<tr><td>San Gregorio, CA</td><td>June 14th</td><td>September 29th</td><td>2</td></tr>
<tr><td rowspan="3">2023</td><td>Davis, CA</td><td>May 24th</td><td>September 12th</td><td>3</td><td>Commercial cultivars: Anasazi, Black Nightfall, Orca, Raven, UCSW Gold, UCSWRed<br>Phenotypic Class 1: BxO-070, OxB-178, BxO-098<br>Phenotypic Class 2: BxO-119, OxB-086, BxO-144<br>Phenotypic Class 3: BxO-139, OxB-175, OxB-094<br>Phenotypic Class 4: BxO-049, BxO-040, BxO-145<br>Phenotypic Class 5: BxO-100, BxO-159, OxB-140<br>Phenotypic Class 6: BxO-013, OxB-139, OxB-179</td></tr>
<tr><td>Tulelake, CA</td><td>May 26th</td><td>September 20th</td><td>2 or 3*</td><td>Commercial cultivars: Anasazi, Black Nightfall, CELRK, Orca, Othello, Raven, UCSW Gold*, UCSWRed*, Viva<br>Phenotypic Class 1: BxO-070, OxB-178, BxO-098<br>Phenotypic Class 2: BxO-119, OxB-086, BxO-144<br>Phenotypic Class 3: BxO-139, OxB-175, OxB-094<br>Phenotypic Class 4: BxO-049, BxO-040, BxO-145<br>Phenotypic Class 5: BxO-100, BxO-159, OxB-140<br>Phenotypic Class 6: BxO-013, OxB-139, OxB-179<br>Tulelake Border: UCSW Gold</td></tr>
<tr><td>Santa Cruz, CA</td><td>June 6th</td><td>October 11th</td><td>2 or 3*</td><td>Commercial cultivars: Anasazi, Black Nightfall*, Orca*, Raven, UCSW Gold*, and UCSW Red*<br>Phenotypic Class 1: BxO-070, OxB-178<br>Phenotypic Class 2: BxO-119, OxB-086<br>Phenotypic Class 3: BxO-139, OxB-175<br>Phenotypic Class 4: BxO-049, BxO-027<br>Phenotypic Class 5: BxO-100, BxO-159<br>Phenotypic Class 6: BxO-013, OxB-139</td></tr>
<tr><td rowspan="2">2024</td><td>Davis, CA</td><td>May 23rd</td><td>September 30th</td><td>2* or 3</td><td rowspan="2">Commercial cultivars: Anasazi*, Black Nightfall*, CELRK*, Orca*, Othello*, Raven*, UCSW Gold*, UCSWRed*, Viva*<br>Phenotypic Class 1: BxO-070, BXO-098, OXB-178<br>Phenotypic Class 2: BXO-119, BXO-144, OXB-086<br>Phenotypic Class 3: BXO-139, OXB-094, OxB-175<br>Phenotypic Class 4: BXO-040, BXO-049, BXO-145<br>Phenotypic Class 5: BXO-100, BXO-159, OXB-140<br>Phenotypic Class 6: BXO-013, OXB-139, OXB-179<br>Tulelake Border: UCSW Gold</td></tr>
<tr><td>Tulelake, CA</td><td>May 31st</td><td>October 3rd</td><td>3 or 4*</td></tr>
</table>

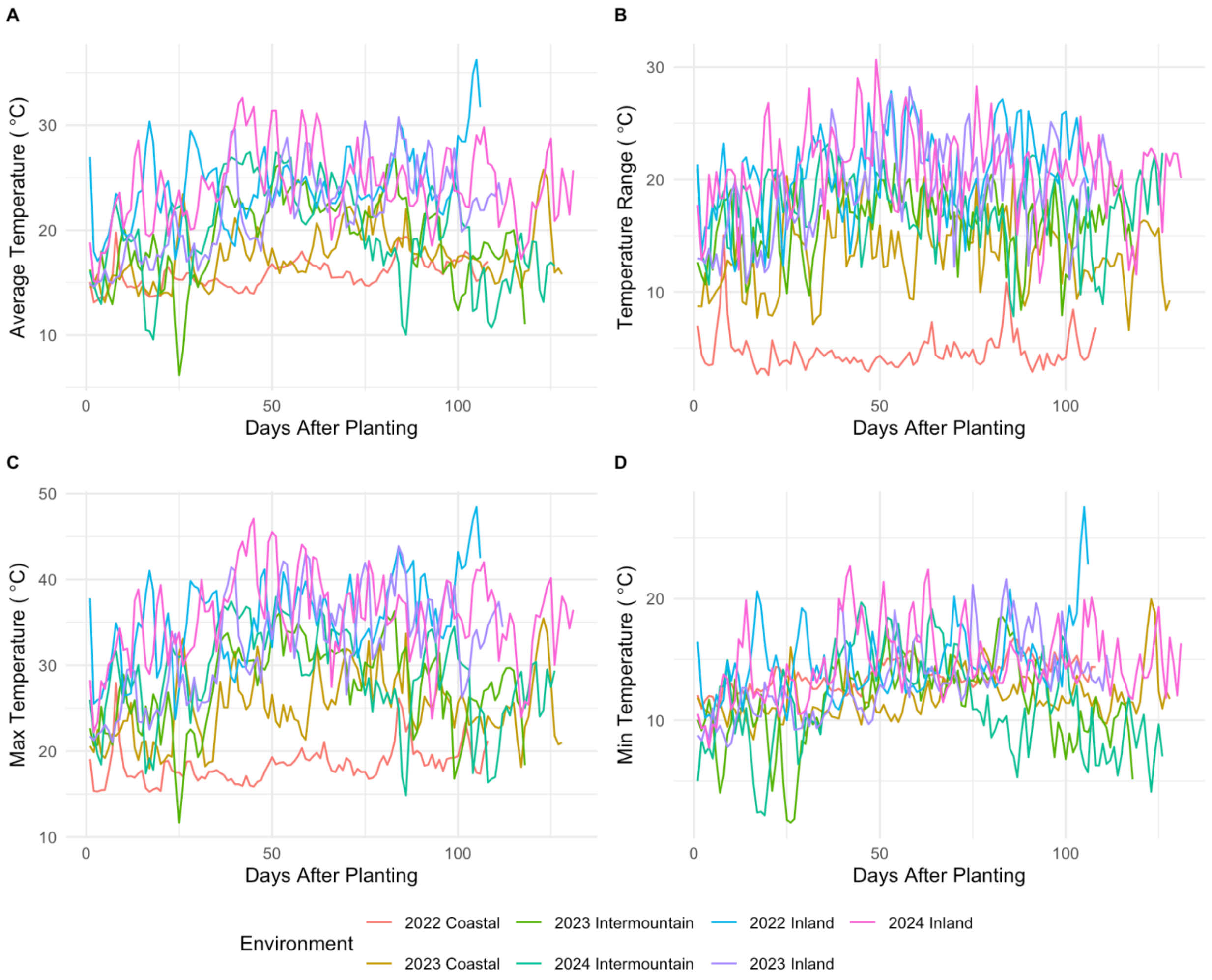


Figure S1. Weather data for each growing environment for the duration of the growing season from planting to harvest.

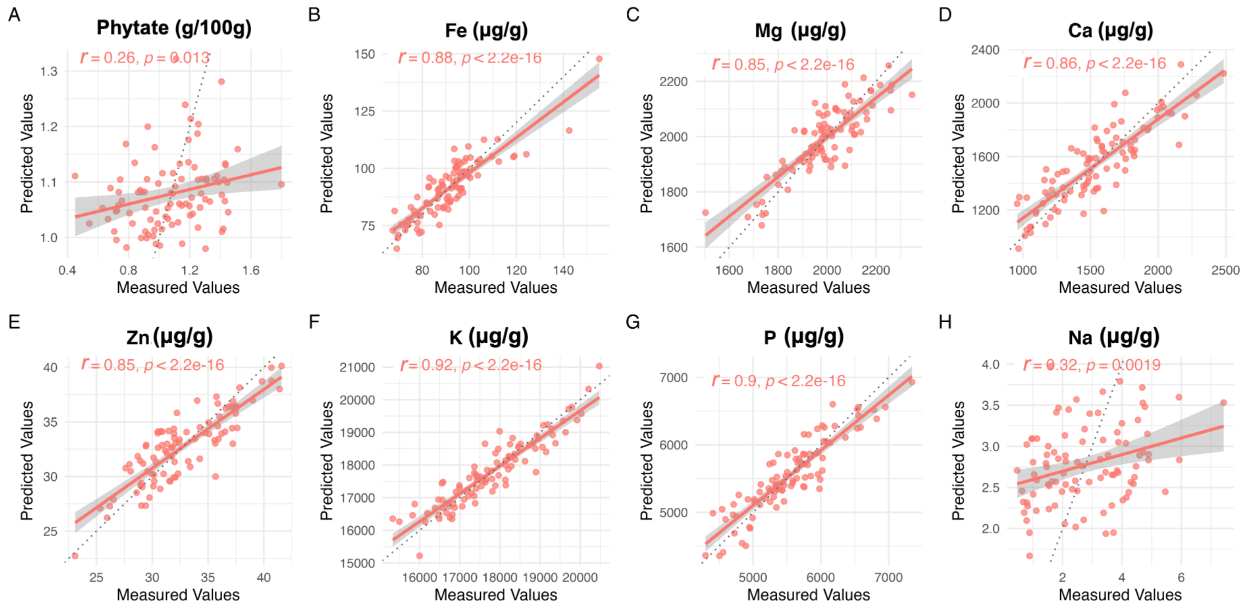


Figure S2. Predicted values for compounds not previously included in the NIRS VPM calibration: A) phytate, B) iron, C) magnesium, D) calcium, E) zinc, F) potassium, G) phosphorus, and H) sodium.

Table S2. By-plot samples scanned via NIRS prior to pooling. Averages and standard deviations for samples from three replicate field plots for Davis and two replicate field plots for San Gregorio.

| Genotype | Location | Protein (%) | Starch (%) | Fat (%) | Crude Fiber (%) | Ash (%) | Moisture (%) |
|---|---|---|---|---|---|---|---|
| BxO-030-PH1 | Davis | 24.44 ± 1.61 | 20.43 ± 1.43 | 2.50 ± 0.49 | 9.66 ± 3.71 | 4.03 ± 0.27 | 8.54 ± 0.70 |
| BxO-049-PH4 | Davis | 25.72 ± 0.86 | 19.24 ± 0.59 | 3.04 ± 0.18 | 6.41 ± 0.17 | 4.17 ± 0.16 | 9.00 ± 0.57 |
| BxO-158-PH2 | Davis | 25.53 ± 0.90 | 19.13 ± 0.98 | 2.12 ± 0.20 | 5.43 ± 0.95 | 3.86 ± 0.19 | 9.37 ± 1.02 |
| OxB-098-PH1 | Davis | 26.14 ± 0.61 | 19.77 ± 0.44 | 2.54 ± 0.14 | 7.41 ± 1.03 | 3.72 ± 0.09 | 7.69 ± 0.40 |
| SouthwestGold | Davis | 25.04 ± 0.42 | 19.94 ± 0.40 | 3.30 ± 0.22 | 7.36 ± 0.46 | 3.89 ± 0.09 | 8.20 ± 0.55 |
| SouthwestRed | Davis | 22.17 ± 0.10 | 22.65 ± 0.24 | 2.34 ± 0.09 | 13.61 ± 0.82 | 4.08 ± 0.14 | 8.45 ± 0.13 |
| Viva | Davis | 24.09 ± 0.75 | 20.19 ± 0.31 | 2.05 ± 0.14 | 14.15 ± 1.48 | 4.16 ± 0.08 | 8.37 ± 0.21 |
| BxO-049-PH4 | San Gregorio | 22.42 ± 0.42 | 22.12 ± 1.07 | 2.09 ± 0.11 | 13.23 ± 1.24 | 4.31 ± 0.47 | 8.71 ± 0.49 |
| BxO-158-PH2 | San Gregorio | 20.49 ± 1.58 | 19.81 ± 1.61 | 3.70 ± 0.28 | 4.15 ± 0.43 | 5.44 ± 0.16 | 13.69 ± 0.52 |
| OxB-098-PH1 | San Gregorio | 20.76 ± 1.69 | 18.25 ± 1.24 | 4.20 ± 0.06 | 3.92 ± 0.11 | 5.29 ± 0.10 | 13.57 ± 0.35 |
| SouthwestGold | San Gregorio | 17.74 ± 1.45 | 21.35 ± 0.81 | 4.78 ± 0.09 | 6.32 ± 2.28 | 6.90 ± 1.03 | 12.50 ± 0.43 |
| SouthwestRed | San Gregorio | 19.05 ± 0.55 | 22.60 ± 1.34 | 3.22 ± 0.17 | 7.39 ± 0.18 | 5.35 ± 0.34 | 10.99 ± 0.46 |
| Viva | San Gregorio | 17.64 ± 1.63 | 22.65 ± 0.85 | 2.97 ± 0.01 | 6.66 ± 0.82 | 5.35 ± 0.01 | 10.79 ± 0.14 |

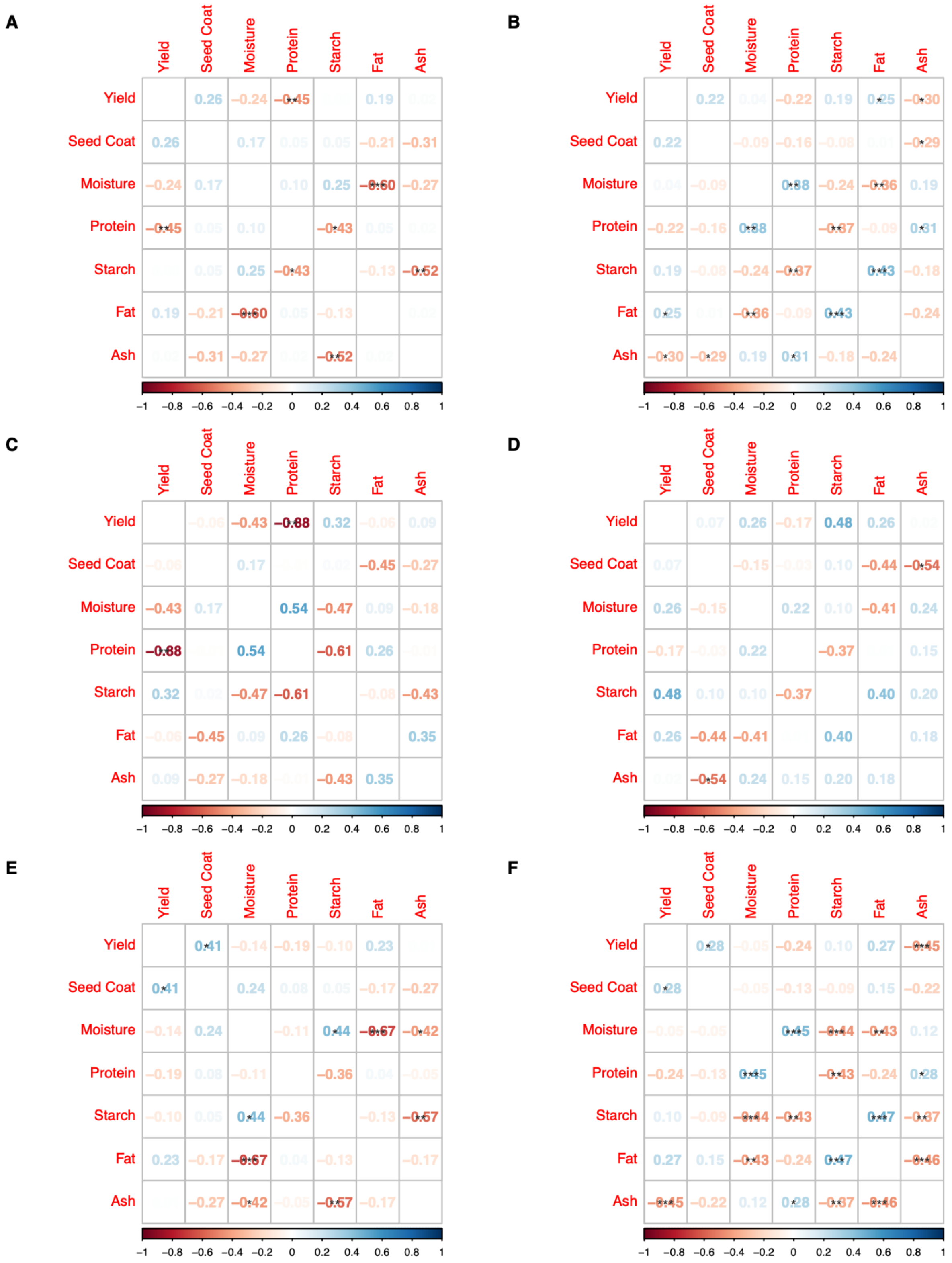

Figure S3. Trait correlations for yield, seed coat, and macronutrients for: A) all field plots in San Gregorio 2022 (n = 32), B) all field plots in Davis 2024 (n = 66), C) commercial cultivar field plots in San Gregorio 2022 (n = 9), D) commercial cultivar field plots in Davis 2024 (n = 16), E)

RIL and parent cultivar (Black Nightfall and Orca) field plots in San Gregorio 2022 (n = 26), and F) RIL and parent cultivar (Black Nightfall and Orca) field plots in Davis 2024 (n = 52). The red-blue color scale represents the *r* value, and significance symbols “***”, “**”, and “*” represent *P*-values of ≤ 0.001, ≤ 0.01, and ≤ 0.05, respectively.
